# Divergent functions of two clades of flavodoxin in diatoms mitigate oxidative stress and iron limitation

**DOI:** 10.1101/2022.10.18.512804

**Authors:** Shiri Graff van Creveld, Sacha N. Coesel, Stephen Blaskowski, Ryan D. Groussman, Megan J. Schatz, E. Virginia Armbrust

## Abstract

Phytoplankton rely on diverse mechanisms to adapt to the decreased iron bioavailability and oxidative stress-inducing conditions of today’s oxygenated oceans, including replacement of the iron-requiring ferredoxin electron shuttle protein with a less-efficient iron-free flavodoxin under iron limiting conditions. And yet, diatoms transcribe flavodoxins in high-iron regions in contrast to other phytoplankton. Here, we show that the two clades of flavodoxins present within diatoms exhibit a functional divergence, with only clade II flavodoxins displaying the canonical role in adaptation to iron limitation. We created CRISPR/Cas9 knock-outs of the clade I flavodoxin from the model diatom *Thalassiosira pseudonana* and found these cell lines are hypersensitive to oxidative stress, while maintaining a wild-type response to iron limitation. Within natural diatom communities, clade I flavodoxin transcript abundance is regulated over the diel cycle rather than in response to iron availability, whereas clade II transcript abundances increase either in iron-limiting regions or under artificially induced iron-limitation. The observed functional specialization of two flavodoxin variants within diatoms reiterates two major stressors associated with contemporary oceans and illustrates diatom strategies to flourish in diverse aquatic ecosystems.

## Introduction

All of life depends on redox-based metabolic pathways driven by the transfer of electrons between protein donors and acceptors. One of the abundant electron shuttles in phototrophs is ferredoxin (Fd), a small, soluble iron-sulfur-cluster containing protein that accepts electrons from the stromal surface of photosystem I during oxygenic photosynthesis and facilitates transfer to acceptors involved in diverse metabolic processes (Mondal and Bruce, 2018). Proteins such as Fds with iron-sulfur clusters are ancient biocatalysts hypothesized to have arisen when oxygen was largely absent, and ferrous iron and sulfide were plentiful (Cammack, 1982). During the great oxidation event approximately 2.4 billion years ago, ferrous iron became oxidized to the ferric form and rapidly precipitated either as ferric hydroxide or as insoluble complexes with anionic salts. Microbial growth in aerobic habitats thus became limited by iron bioavailability (Imlay, 2006), as found in about a third of today’s oceans (Behrenfeld and Milligan, 2013; Boyd et al., 2007; Moore et al., 2013). Moreover, iron-sulfur-containing proteins are particularly sensitive to damage via oxygen and reactive oxygen species (ROS). And yet, contemporary marine phytoplankton (cyanobacteria and photosynthetic eukaryotes) continue to rely on Fd and may sequester up to 30–40% of their cellular iron within these proteins (Erdner, 1997). Reliance on Fd for electron shuttling during oxygenic photosynthesis thus carries with it both a requirement for sufficient iron bioavailability and an enhanced sensitivity to the ROS generated during photosynthesis.

Under iron limiting conditions, photosynthetic microbes commonly replace the iron-containing Fd with flavodoxin, a functional homologue (Smillie, 1965; Hutber et al., 1977; Sandmann et al., 1990). Flavodoxin uses a flavin mononucleotide (FMN) as the co-factor and is a less efficient electron shuttle, that nonetheless allows continued photosynthetic electron transport when iron is scarce (WG Zumft and Spiller, 1971; Yoch and Valentine, 1972). The absence of flavodoxin from land plants and certain coastal algal species led to the premise that flavodoxin is lost from species in iron-rich environments, including terrestrial systems (Pierella Karlusich et al., 2015). However, the downregulation of Fd in response to adverse conditions (Singh et al., 2010, 2004) and the induction of flavodoxin in the cyanobacteria *Synechocystis* under oxidative stress (Jeanjean et al., 2003; Kojima et al., 2006; Singh et al., 2004) suggests additional physiological roles for flavodoxin within phytoplankton.

Expression levels of flavodoxin relative to Fd (inferred either from transcript or protein levels) have been used in several studies to evaluate whether natural communities of marine phytoplankton experience iron limitation *in situ* (Boyd et al., 1999; DiTullio et al., 2005; Erdner and Anderson, 1999; Jones, 1988). Metatranscriptome studies indicate that, as expected, most major eukaryotic phytoplankton lineages (chlorophytes, haptophytes and dinoflagellates) regulate flavodoxin transcript abundances in response to iron availability, with reduced flavodoxin transcript levels in iron-replete environments (Caputi et al., 2019; Carradec et al., 2018). A closer examination of environmental metatranscriptomes (Marchetti et al., 2012; Carradec et al., 2018; Caputi et al., 2019) finds that diatom flavodoxin transcript abundances are not inversely correlated with iron availability as expected, and flavodoxins are often detected at high abundances under iron replete conditions. Although direct comparisons between environmental flavodoxin and Fd transcript abundances are complicated by the fact that Fd is encoded on the plastid genomes of some diatoms (Groussman et al., 2015; Gueneau et al., 1998; Lommer et al., 2010; Oudot-Le Secq et al., 2007; Roy et al., 2020), diatoms appear to regulate the transcription of flavodoxin differently compared to other phytoplankton. A potential explanation for the distinctive flavodoxin transcriptional patterns of natural communities of diatoms comes from the discovery in a few model diatoms that flavodoxin diverged into two distinct phylogenetic clades with only those flavodoxins from clade II induced upon iron limitation, suggesting that clade I flavodoxins may play a different role within these species (Whitney et al., 2011).

Here, we combine genetic surveys of environmental and experimental datasets with gene knockout strategies in a model diatom to elucidate the differentiation of clade I and clade II flavodoxins in phytoplankton and their potential roles in diatoms. Our results from both model diatoms and natural phytoplankton communities indicate that these two flavodoxin clades represent a functional divergence where the clade II flavodoxins in stramenopiles play a role in adaptation to low iron conditions and the clade I variant likely conveys adaptation to oxidative stresses. This functional divergence likely augments the ability of diatoms to flourish in diverse ecosystems.

## Results

### Clade I flavodoxins emerged within a subset of stramenopiles

The original description of clade I and II flavodoxins was based on sequences from 6 model diatoms and one non-diatom stramenopile (Whitney et al., 2011), with the potential for functional distinction derived from transcriptional responses of two *Thalassiosira* diatoms, each encoding a different clade of flavodoxin. We supplemented this work by examining publicly available transcript data and found that in the three model diatoms that encode both clade I and clade II flavodoxins, only the clade II flavodoxins were significantly induced upon iron limitation (Graff van Creveld et al., 2016; Lommer et al., 2012; Mock et al., 2017; Smith et al., 2016) (summed in Table S1), reiterating the potential for different functional roles of these proteins.

To determine the distribution of clade I and clade II flavodoxins in additional marine phytoplankton, we screened publicly available genomes and transcriptomes of over 500 marine protists, bacteria, and archaea (Coesel et al., 2021) for flavodoxin sequences using a custom-made hidden Markov model (hmm)-profile of the flavodoxin domain adapted from PF00258 (e < 0.001; hmmsearch; Eddy, 1998; Finn et al., 2014). The flavodoxin domain was detected in 1191 flavodoxin genes and included 332 genes that clustered with known photosynthetic flavodoxin proteins; the remaining sequences displayed similarity to a diverse and distant clade of non-photosynthesis-related proteins (Fig. S1A, supplemental fasta file 2). The presumed photosynthesis-associated flavodoxins segregate into four clades (Fig. 1A, Fig. S1A). Two clades were composed of photosynthetic flavodoxins from green algae, one that grouped with gamma-proteobacteria (highlighted in pink, Fig. 1A, Fig. S1B) and another that grouped with cyanobacteria and dinoflagellates (highlighted in green, Fig. 1A, Fig. S1B). Within these clusters, the labelled *Synechococcus* and *Prochlorococcus* flavodoxins were previously shown to be induced in response to iron limitation (Kashtan et al., 2014; Mackey et al., 2015; Thompson et al., 2011; Yousef et al., 2003). One iron limitation-induced *Synechococcus* flavodoxin grouped within the clade of green algae and gamma-proteobacteria flavodoxins (highlighted in pink, Fig. 1A, Fig. S1B); all other iron limitation-induced *Synechococcus* flavodoxins clustered within the green algae clade that contained proteins from cyanobacteria and dinoflagellates (highlighted in green, Fig. 1A, Fig. S1B).

**Figure 1.**
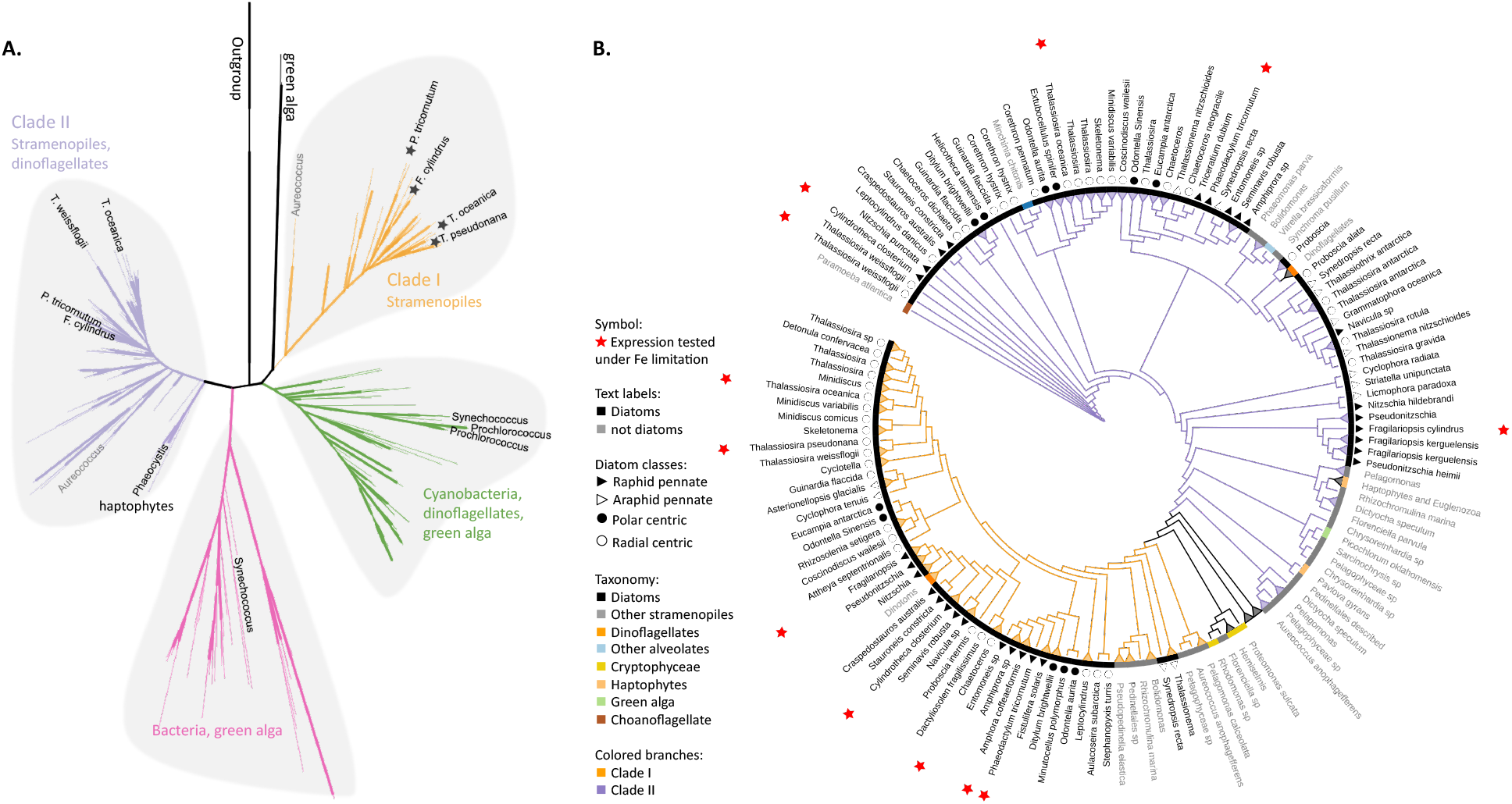
Phylogeny of flavodoxins in marine microorganisms. **A.** Maximum-likelihood (RAxML) phylogenetic tree of photosynthetic flavodoxins (see Fig. S1A for the full tree, Fig. S1B for taxonomic annotation); non-photosynthetic flavodoxins as well as flavodoxin domains in other proteins are collapsed as an “outgroup”. Thick branch lines represent bootstrap support greater than 0.7. Different clades are marked with colors: clade I (orange), clade II (purple), bacteria/green algae clade (pink), cyanobacteria/ dinoflagellate/green alga clade (green). Labeled taxa indicated with black labels were experimentally tested for response to iron limitation; tested flavodoxins indicated with black starts were not induced (Tables S1, S2). *Aureococcus* (indicated in gray) has not been experimentally tested in response to iron limitation. **B.** Unrooted maximum-likelihood (RAxML) phylogenetic tree of clade I and II flavodoxins with 56 additional stramenopile strains (listed in Table S2). Thick branch lines represent bootstrap support greater than 0.7. Branches collapsed (indicated with triangles at tips) for non-stramenopiles and at genus level within stramenopiles (see Fig. S1C for the full tree). Branch colors represent clade (I, II, orange and purple respectively). Colored strip represents taxonomy, dinoflagellates with diatom endosymbionts are labeled ‘dinotoms’. Red stars represent flavodoxins with expression experimentally tested under iron limitation.

Clade I and II flavodoxins, as originally identified by Whitney (2011), were detected exclusively in eukaryotic–algae originally derived from a secondary endosymbiosis of a red alga (Fig. 1A, Fig. S1B). Clade I flavodoxins were detected in stramenopiles (highlighted in orange, Fig. 1, Fig. S1B) and two dinoflagellates (*Durinskia baltica* and *Kryptoperidinium foliaceum*). Both dinoflagellates express proteins originating from their diatom endosymbionts (Hehenberger et al., 2016; Imanian et al., 2010; Imanian and Keeling, 2007; Yamada et al., 2019). The flavodoxin from the stramenopile *Aureococcus anophagefferens*, previously defined as a clade I flavodoxin (Whitney et al., 2011), groups with another unclassified pelagophyte (Fig. 1A, Fig. S1B). This branch was defined here as the base of clade I. None of the previously tested clade I flavodoxin genes transcriptionally respond to iron limiting growth conditions (stars, Fig. 1A, Table S1). Clade II flavodoxins are phylogenetically more diverse than clade I flavodoxins, and consist of sequences from diatoms, non-diatom stramenopiles, haptophytes and dinoflagellates (*Karenia brevis* and *Karlodinium micrum*) (Fig. 1A, Fig. S1B). The base of clade II is defined here by a cluster of haptophyte sequences that includes a *Phaeocystis* flavodoxin (labeled) experimentally induced under iron limitation (Wu et al., 2019). Known diatom iron-limitation responsive flavodoxins all cluster within clade II (Table S1), consistent with a functional divergence between clade I and clade II flavodoxins within the stramenopiles.

To determine the distribution of clade I and clade II flavodoxins within the diverse stramenopile lineage more thoroughly, we queried 56 additional publicly available stramenopile transcriptomes and genomes (Table S2, supplemental fasta file 3). Flavodoxins from both clades appear to be similarly distributed amongst open ocean and coastal isolates, with no evidence that clade II flavodoxins are retained only in species isolated from low-iron environments (Fig. S1B, Table S2). Diatom clade I flavodoxins from pennate and centric diatoms form a tight cluster with high bootstrap support (Fig. 1B, Fig. S1C), with multiple copies detected in a subset of species (Table S2, Fig. S1D). Many examined taxa encode both clade I and clade II flavodoxins. About half of the taxa that appear to encode a flavodoxin from only one clade (Fig. S1D), although this absence may reflect culture conditions rather than absence from their respective genomes as most available sequence data are derived from transcriptomes (Table S2). Nonetheless, the absence of clade I flavodoxin sequences appears more common in stramenopiles that are more distantly related to diatoms, such as *Phaeomonas* (Pinguiophyceae; Fig. S1E), while stramenopiles more closely related to diatoms, such as members of Pelagophyceae and Dictyochophyceae, commonly encode both clade I and clade II flavodoxins (Fig. 1B, Fig. S1E, Table S2).

To explore a potential molecular basis for the clade I and II protein divergence, we aligned stramenopile flavodoxins to the flavodoxin sequence of the red alga *Chondrus crispus* (supplemental fasta file 4), as its structure is well studied and the FMN binding sites were previously identified (Fukuyama et al., 1992). As expected, amino acid side chains that form hydrogen bonds with the FMN (underlined in Fig. S1F) are conserved between the two clades. The clade I and clade II flavodoxin sequences differ from each other at amino acids 57 and 103 (numbered according to *C. crispus* sequence, Fig. S1F). At amino acid 57, the dominant amino acid is asparagine (N) in *C. crispus* and clade I proteins, and histidine (H) in clade II proteins. At amino acid 103, the dominant amino acid is cysteine (C) in *C. crispus* and clade II proteins, and alanine (A) in clade I proteins. Both distinguishing amino acid changes occur near the FMN-binding tryptophan and tyrosine (W56, Y98, indicated with asterisks in Fig. S1F), consistent with distinct functions for the two flavodoxins.

### Diatom flavodoxins transcriptional responses to oxidative and iron stress

Given the observed dual role of the cyanobacterial flavodoxin in adaptation to iron limitation and oxidative stress (Jeanjean et al., 2003; Kojima et al., 2006; Singh et al., 2004), we hypothesized that the two clades in stramenopiles represented a functional divergence with the iron limitation response specific to clade II, and the oxidative stress response specific to clade I flavodoxins. To test this hypothesis, we examined transcriptional responses of 5 diatom species exposed to either H_2_O_2_-induced oxidative stress, or to iron limitation. We chose two closely related model centric diatoms: the estuarian-isolated *Thalassiosira pseudonana* and the open ocean-isolated *Thalassiosira oceanica*, as well as three non-model open ocean-isolated diatoms: two pennates *Amphora coffeaeformis* and *Cylindrotheca closterium* and one centric *Chaetoceros sp*., none of which had publicly available genome or transcriptome sequences. Each diatom was grown in either iron-replete or iron-limiting conditions for 3-6 days until the low-iron cultures displayed a decrease in maximum photochemical yield of photosystem II (Fv/Fm) (Fig. S2A-C, Fig. 2A, Table S3), indicative of iron limitation, at which point transcriptome samples were collected for both the iron-limited and iron-replete conditions. A subset of iron-replete cultures was exposed to oxidative stress, mimicked by a lethal dose of H_2_O_2_, and transcriptome samples were collected about 1.5 h after exposure, when the cell phenotype was unaltered. The three conditions elicited distinct overall transcriptional responses in the 5 diatom species as seen in multidimensional scaling (MDS) plots (Fig. S2D).

**Figure 2.**
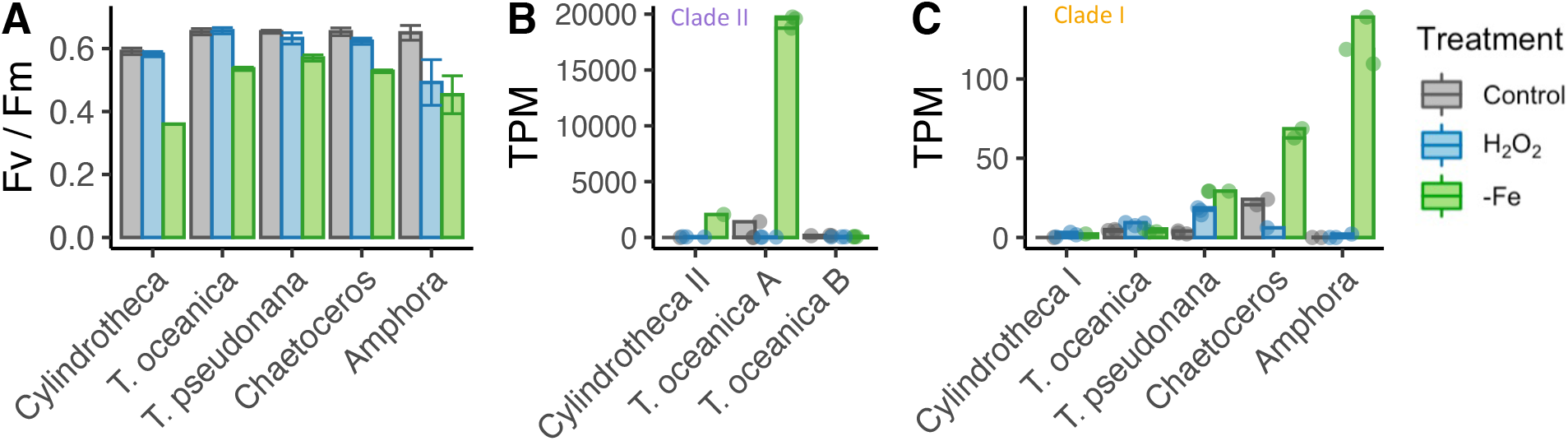
Iron limitation and oxidative stress in diatom cultures. **A.** Photosynthetic efficiency (Fv/Fm) of five diatom species, before harvesting the cells for each transcriptome, Error bars represent the standard deviation of biological triplicates. Clade II (**B**) and clade I (**C**) flavodoxins expression in response to iron limitation and H_2_O_2_ treatment. *T. oceanica* clade II flavodoxins were previously named FLDA1 and FLDA2, marked here as A and B accordingly. *Cylindrotheca* encodes one flavodoxin from each clade, marked here I and II according to the clade. Note the different Y axis scales in panels B and C. Individual measurements are marked in circles, maximal values in colored bars.

The distribution and transcriptional patterns of clade I and clade II flavodoxin genes differed within the five examined diatom species. *Cylindrotheca* and *T. oceanica* transcribed both clade I and II flavodoxins; *T. oceanica* transcribed two isoforms of clade II flavodoxins (labeled here A, B, Fig. 2B); *T. pseudonana, Chaetoceros*, and *Amphora* transcribed only clade I flavodoxins. The transcription levels of clade II genes from *T. oceanica* (clade II-A) and *Cylindrotheca* were significantly elevated in response to iron limitation (5.3 and 9.5-fold change, respectively, FDR < 0.01, Fig. 2B), whereas the clade I genes in these diatoms did not display a significant change in transcript abundance either after exposure to H_2_O_2_ or under iron limitation (FDR = 1, Fig. 2C). The *T. oceanica* clade II-B gene transcript levels were not responsive to either treatment (Fig. 1B), in agreement with previous results (Table S1, Lommer et al., 2012). The transcribed clade I flavodoxins of *T. pseudonana*, *Chaetoceros*, and *Amphora* were elevated under iron limitation (3.2, 1.5, 9.9-fold change, respectively, FDR < 0.01, Fig. 2C). Suggesting that clade I flavodoxins are induced in iron limitation only when clade II flavodoxins are absent. Only the *T. pseudonana* clade I flavodoxin gene was significantly differentially transcribed in response to H_2_O_2_ treatment (2.4-fold increase, FDR < 0.01, Fig. 2C). Together these results confirm a role for clade II flavodoxins in response to iron limitation and suggest that the role of the clade I flavodoxin within a species may depend on whether clade II flavodoxin is also transcribed.

### Functional role of a clade I flavodoxin

To disambiguate the potential for functional redundancy or synergy between clade I and II flavodoxins within a species, we generated a clade I flavodoxin gene knock out (KO) in *T. pseudonana*, as this diatom encodes a single copy of the clade I gene (TpFlav, Thaps_19141) and lacks a clade II gene. CRISPR/Cas9 was used to generate three independent KO lines, each with deleted FMN binding sites (Fig. S3A-C). Two wild type-like lines (WT) were also retained that were transformed with the same plasmid, but in which the flavodoxin gene was not edited (Fig. S3B, C). Cell growth, Fv/Fm, and the final carrying capacity were not significantly different between WT and KO cells under iron-replete growth conditions (Fig. 3A, B, Fig. S3D, E). Cell counts were similar between WT and KO lines, reaching about 3 million cells per ml at day 12 (WT 3.04 ±0.25, KO 2.93 ±0.11, Fig. 3A, Fig. S3D). Photosynthetic efficiency was also similar between WT and KO cells (0.64 ±0.016, 0.63 ±0.040 respectively, at day 1, Fig. 3B, Fig. S3E). The WT and KO lines displayed comparable reductions in cell division and Fv/Fm under iron limitation and after ~3 days, nearly reached a maximum of about 400,000 cells per ml (Fig. 3A, Fig. S3F), and decreased in Fv/Fm from 0.63 ±0.016 (all cell lines in replete conditions) to 0.57 ±0.016 (KO lines) or 0.57 ±0.004 (WT lines) (Fig. 3B, Fig. S3G). Thus, both wild-type and KO *T. pseudonana* lines respond poorly to iron limiting conditions, which suggests that wild type cells do not replace their iron-requiring Fd with flavodoxin during adaption to low iron. A difference in phenotype between WT and KO cell lines emerged, however, after treatment with H_2_O_2_ (Fig. 3C). Exposure to 100 μM H_2_O_2_ killed ~73% of WT cells after 24 h. In contrast, exposure of KO cells to the same dose of H_2_O_2_ resulted in the death of ~100% of the KO cells after 24 h, comparable to what was seen when WT cells were exposed to twice the amount of H_2_O_2_ (200 μM). The hyper-sensitivity of KO lines to H_2_O_2_ (Fig. 3C, Fig. S3H) indicates that clade I flavodoxins play a role in the oxidative stress response.

**Figure 3.**
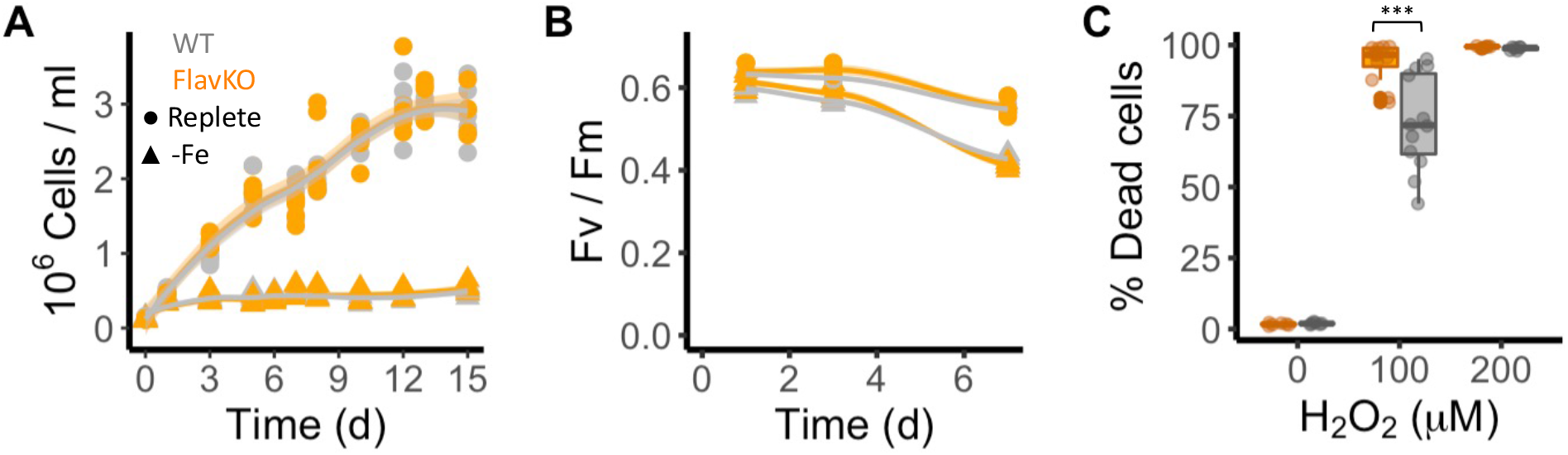
Response of *T. pseudonana* flavodoxin KO and WT lines to iron limitation and oxidative stress. Three independent WT (gray) and flavodoxin KO (orange) lines were grown under iron-replete (circles) and iron-limiting (triangles) conditions for several days. **A.** Cell abundance, measured by flow cytometry. **B.** Photosynthetic efficiency, measured by phytoPAM. Individual measurements marked in symbols, means of triplicates in lines. **C.** Percentage of Sytox Green-positive (dead) cells, measured by flow cytometry 24 h after treatment with 0, 100 or 200 μM H_2_O_2_. Box plots combine two independent experiments, ANOVA test between WT and KO (indicated with ***), significant 9•10^-5^.

### Environmental flavodoxin transcription in the North Pacific Ocean

To expand our analysis beyond laboratory studies, and explore the potential roles of clade I and clade II flavodoxins in natural communities, we interrogated a total of 120 eukaryotic surface-water metatranscriptomes sampled during three research cruises in the North Pacific Ocean: Gradients 1 cruise (April/May 2016) transited south/north along ~158 °W, from 21 °N to 38 °N (10 stations, triplicates); Gradients 2 cruise (May/June 2017) transited south/north along ~158 °W, from 24 °N to 42.5 °N (10 stations, triplicates), and a diel sampling cruise (July/August 2015) was conducted in the North Pacific Subtropical Gyre (NPSG), at ~156.5° W, 24.5° N near station ALOHA (24 times points, duplicates); an on-deck nutrient-addition experiment was conducted on Gradients 2 at 37 °N, 158 °W (incubations: 4 conditions, triplicates) (Fig. 4A). Clade and taxonomic affiliations at either the species or genus level were assigned to each assembled environmental transcript (contig) based on their phylogenetic placement (Barbera, 2016; Barbera et al., 2019) within our stramenopile-expanded photosynthetic flavodoxin reference tree (Fig. S4).

**Figure 4.**
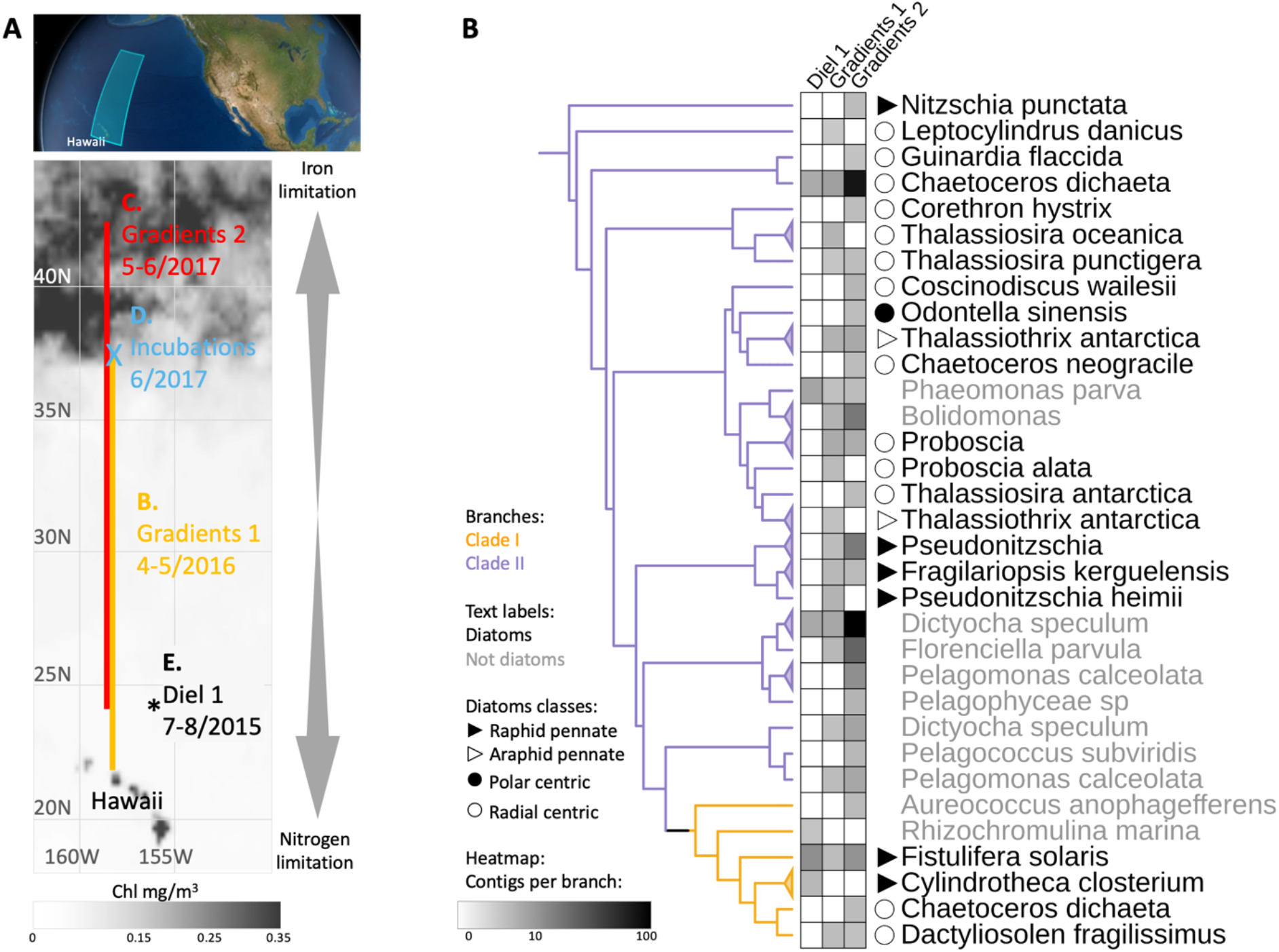
Detection of the two flavodoxin clades in the North Pacific. **A.** Overview of sampling area with a background of satellite, average chlorophyl estimate, from May 9 to June 26, 2017. Data provided by the Ocean Colour Thematic Center at the Copernicus Marine environment monitoring service (CMEMS) and visualized with SimonsCMAP (Ashkezari et al., 2021). Cruise dates (month/year) and locations and the site of the on-deck incubation during Gradients 2 are marked. **B.** Heatmap representing the number of stramenopiles clade I or II flavodoxin contigs detected in each cruise placed on the flavodoxin reference phylogenetic tree. Branches consisting of species (and strains) from the same genus were collapsed and marked with triangles at the edges and labeled according to the highest taxonomic rank. The un-collapsed clade I and clade II region of the reference tree, with environmental placements is shown in Fig. S4. Branch colors represent clade I (orange) and clade II (purple). The genus and species names are in black for diatoms, or gray for other stramenopiles.

Contigs for the two flavodoxin clades were detected in metatranscriptomes from all three expeditions, with a greater number and phylogenetic diversity of clade II flavodoxin contigs, particularly along the Gradients 1 and 2 transects (Fig. 4B). The majority of environmental clade II flavodoxins were most closely related to the reference sequence of the radial centric diatom *Chaetoceros dichaeta*, originally isolated from the South Atlantic; additional clade II contigs were distributed amongst 11 other diatom genera (Fig. 4B, Fig. S4). Within the non-diatom stramenopiles, a majority of clade II flavodoxin contigs were assigned to *Dictyocha* and *Florenciella*, with additional contigs distributed amongst 5 other genera (Fig. 4B, Fig. S4). The clade I flavodoxins were assigned primarily to *Fistulifera solaris*, a small pennate diatom similar to those dominating the diatom population in this area (Dore et al., 2008; Villareal et al., 2012); additional clade I contigs were assigned to three other reference diatoms, including *Chaetoceros dichaeta*, and two non-diatom stramenopiles (Fig. 4B, Fig. S4).

Community composition, nutrient availability (Gradoville et al., 2020; Juranek et al., 2020; Lambert et al., 2022; Park et al., 2022; Pinedo-González et al., 2020), and flavodoxin transcriptional patterns (Fig. 5A, B, Fig. S5, Table S4) varied spatially along the Gradients 1 and 2 south/north transects. On Gradients 1, iron (Fe) and nitrate (N) displayed the expected opposing concentration patterns with increasing N and decreasing Fe concentrations north of the NPSG, resulting in low N/Fe (~0.01 μM/nM) within the NPSG and 3-4 orders of magnitude higher N/Fe (~10-100 μM/nM) within the transition zone and further north (Fig. 5C, D). In contrast, on Gradients 2, Fe increased within the transition zone due to increased dust deposition (Pinedo-González et al., 2020) resulting in a relatively low N/Fe (< 10 μM/nM) throughout the transition zone up to ~40 °N (Fig. 5E, F). We compared flavodoxin transcript abundances between genera across the two cruise transects (Fig. 5A, B) by mapping the short sequence reads to our flavodoxin contigs and normalizing these counts to the total transcript concentrations assigned to their respective taxonomic order for each sample (scheme in Fig. S5A). Clade I flavodoxin transcripts were detected for two genera, *Fistulifera* and *Dactyliosolen*, and the relative abundances of these transcripts were higher within the NPSG where N/Fe ratio remained <1 (μM/nM) for both cruises (Fig. 5A-F). In contrast, the clade II flavodoxin transcript abundances displayed spatial differences between the two cruises. On Gradients 1, relative clade II flavodoxin transcript abundances increased within the transition zone, peaking at 33 °N where the N/Fe ratio reached >37 (μM/nM), for all detected genera except *Chaetoceros*, which displayed the highest relative transcript abundances at 37 °N, the most northern station of this transect (Fig. 5A, C, D). Consistent with the higher Fe concentrations detected on Gradients 2 (Park et al., 2022; Pinedo-González et al., 2020), relative clade II transcript abundance did not undergo as great an increase within the transition zone as seen during the Gradients I cruise. Instead, transcript abundances for 4 of 10 detected genera did not peak until 40 °N, the station at which N/Fe first reached >6 μM/nM (Fig. 5B, E, F). These results indicate that clade II flavodoxins are expressed by diverse genera specifically under relatively high-nitrate, low iron conditions, whereas clade I flavodoxin transcription is restricted to a few diatom genera in oligotrophic, non-iron limiting conditions.

**Figure 5.**
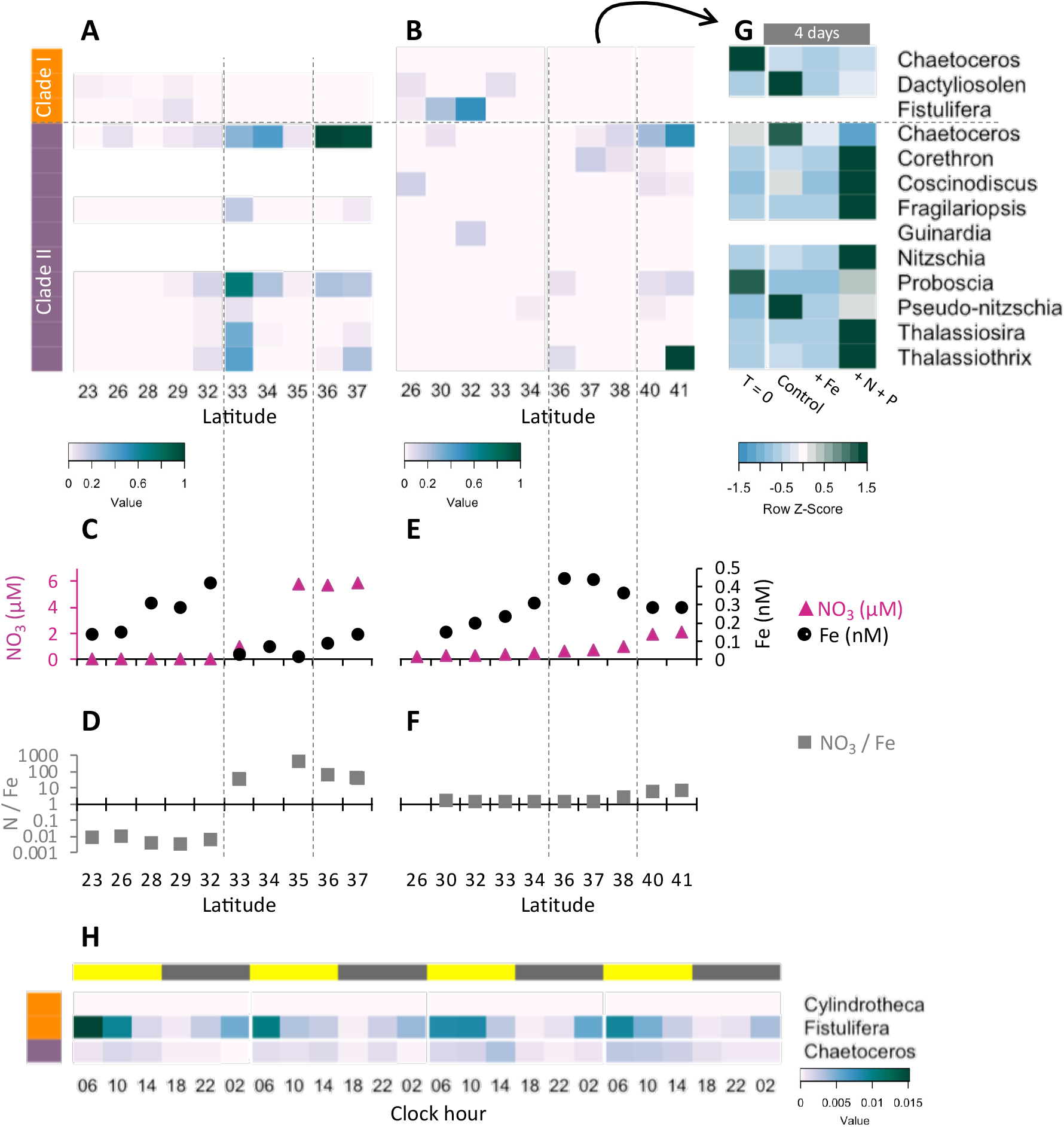
Transcriptional patterns of diatom flavodoxin genes in the North Pacific. **A, B.** Heatmaps of relative diatom flavodoxins transcription across the Gradients 1 transect (**A**) or Gradients 2 transect **(B**). Flavodoxin sequence reads per liter were summed at the genus level and normalized to the cumulative number of reads per liter. White rows indicate where flavodoxin transcripts were not detected. **C, E.** Total dissolved iron (Fe, black circles) and nitrate (NO_3_, pink triangles) concentrations across Gradients 1 (**D**) and Gradients 2 (**F**) transects (Gradoville et al., 2020; Juranek et al., 2020; Park et al., 2022; Pinedo-González et al., 2020). **F, G.** Nitrate to iron ratio (gray rectangles) of these transects. **G.** Flavodoxin transcription following nutrient enrichment incubations conducted at Gradients 2, 37 °N. Water were sampled at T=0 and incubated with no added nutrients (Control) or with 1 nM FeCl_3_ (+Fe), or 5 μM NO_3_ and 0.5 μM PO_4_ (+N +P) and sampled for metatranscriptomes after 4 days. Transcripts per liter are row normalized. **H.** Diatom relative flavodoxin transcript abundance across the diel cycle, sampled during the Diel 1 cruise. Upper bar represents day (yellow) or night (gray). Side bars represent flavodoxin clade I (orange) or clade II (purple). Note color scale differs from those of **A, B, G**.

To determine how North Pacific diatom genera at a given location respond to changes in N/Fe, we conducted on-deck incubation experiments on Gradients 2 using trace-metal clean seawater collected from 37 °N, 158 °W where the *in situ* N/Fe ratio was 0.24 (μM/nM, Fig. 5F). Seawater samples were maintained in on-deck, temperature-controlled incubators for 4 days with triplicate bottles amended with either no nutrient additions (control); 1 nM FeCl_3_ (+Fe), to alleviate potential iron limitation; or 5 μM NO_3_ and 0.5 μM PO_4_ (+N +P), which was expected to enhance iron limitation. Samples for metatranscriptome analysis were taken at T=0 and at T=4 days. For this study, the flavodoxin transcripts per liter were summed for each genus in each treatment and normalized to the total flavodoxin transcripts per genus detected in all treatments (e.g. ‘row-normalized’) (Fig. 5G, Fig. S5A). Flavodoxins from both clades were detected in these experiments. Clade I flavodoxins transcripts were low in both *in situ* samples (T=0) and in the experimental treatments (T=4 days) (37 °N Fig. 5B, Table S4) with both nutrient amendment conditions (Fe or N+P) resulting in a reduction in relative transcript abundances assigned to *Chaetoceros* and *Dactyliosolen* (Fig. 5G). In contrast, relative clade II transcript abundances increased significantly in the N+P amendment treatment compared to both the controls and the Fe amendment in all genera except *Chaetoceros, Proboscia*, and *Pseudo-nitzschia* (Fig. 5G). *Proboscia* displayed the highest relative transcript abundances *in situ* (T=0) and *Chaetoceros* and *Pseudo-nitzschia* displayed the highest relative transcript abundances in the control. Together, these results indicate that diatoms can respond to iron limitation by upregulating transcription of clade II flavodoxins, which supports the inference that the relatively low abundance of clade II transcripts detected *in situ* at 37 °N reflected iron-replete conditions at the time of sampling. Importantly, these experiments also illustrate the underlying diversity within diatom genera in their response to iron availability and highlight additional diatoms such as *Pseudo-nitzschia* and *Chaetoceros* for future laboratory studies of adaptive responses to iron limiting conditions.

To further explore a potential role for clade I flavodoxins, we examined transcriptional patterns of clade I and clade II flavodoxins in metatranscriptomes collected in duplicate every 4 h for 4 days within the NPSG (diel 1 cruise, Wilson et al., 2017), a low nitrogen, high-light environment. Each day the photosynthetically active radiation (PAR) reached ~2000 μmol photons·m^-2^·sec^-1^ (Coesel et al., 2021), conditions expected to enhance oxidative stress. Flavodoxin transcripts per liter were normalized by the total reads assigned to each taxonomic order, as done for the Gradients transects samples (Fig. S5A). Clade II transcript abundances were low throughout the diel cycle (Fig. 5H), in agreement with the low transcript levels detected at similar latitudes along the Gradients transects (~24 °N, Fig. 5A, B) representative of the non-iron limiting conditions at these stations. In contrast, clade I flavodoxins exhibited a clear diel pattern in transcript abundance. Most contigs were assigned to *Fistulifera solaris*, and 13 out of 19 contigs had a significant peak in transcript abundance at 6 AM (RAIN analysis, BH correlation < 0.05) that then decreased gradually throughout the day and reached a minimum at dusk. The increase in clade I transcript abundances during the night beginning at ~10 PM - 2 AM suggests a diel-regulated anticipation of the high light exposure after dawn.

## Discussion

Marine phytoplankton rely on multiple mechanisms to adapt to the oxygen-rich, iron-poor conditions of contemporary oceans. We focused on the role of flavodoxin, an iron-independent electron shuttle protein that can replace the more efficient iron-dependent protein ferredoxin under iron limiting conditions, and which is less sensitive to reactive oxygen species. Both ferredoxin and flavodoxin were presumably present in the cyanobacteria engulfed during the primary endosymbiosis that gave rise to the plastids of the green, red, and glaucophyte algae (Campbell et al., 2019). During the subsequent secondary endosymbiosis of a red alga, these proteins were transferred to stramenopile, dinoflagellate, and haptophyte lineages, that together dominate carbon flux in contemporary global oceans. The complex evolutionary history of flavodoxin has resulted in distributions scattered across all domains of life, a pattern hypothesized to reflect both multiple horizontal gene transfers and gene loss in organisms that evolved in iron-rich environments (Pierella Karlusich et al., 2015; Pierella Karlusich and Carrillo, 2017). Our expanded genetic survey of flavodoxins in marine microorganisms confirmed the expected relatedness of flavodoxins from cyanobacteria and primary endosymbionts (red and green algae), and their divergence from those of secondary endosymbionts. Responsiveness to iron limitation by both marine cyanobacterial and green algal flavodoxins supports the proposition that iron availability is a strong selective pressure on retention of flavodoxin within the genomes of these organisms (Pierella Karlusich et al., 2015). The distribution of the clade I and clade II flavodoxins within secondary (and tertiary) endosymbionts points towards additional selective pressures. Clade II flavodoxins from model organisms display responsiveness to iron limitation, exhibiting the expected canonical replacement of ferredoxin by flavodoxin under iron limiting conditions. And yet, the distribution of clade II flavodoxins across stramenopiles lineages appears unrelated to contemporary iron conditions. Clade I flavodoxins are restricted to stramenopiles, apparently differentiated in the Diatomista after their divergence from the Chrysista (Fig. S1E). Clade I flavodoxins appear to have been retained across diverse diatoms and are often present in multiple copies, regardless of whether the species was originally isolated from the iron-poor open ocean or the iron-rich coastal environment suggesting alternative selective pressures.

Prior to our study, relatively little attention was given to the potential role of flavodoxin as part of an oxidative stress response in marine phytoplankton. However, in *Escherichia coli*,overexpression of endogenous flavodoxin resulted in enhanced resistance to oxidative stress (Zheng et al., 1999), and ectopic expression of cyanobacterial flavodoxin in land plants leads to enhanced tolerance to a broad range of stresses, including oxidative stress (Blanco et al., 2011; Ceccoli et al., 2011; Mayta et al., 2019; Tognetti et al., 2006; Zurbriggen et al., 2008). Thus, when expressed at high levels, the known iron-responsive flavodoxins can mitigate additional stresses that may compromise Fd functionality. Our laboratory studies with five diatom species confirmed that clade II flavodoxin expression is an adaptation to iron limitation, presumably resulting in the replacement of Fd as an electron shuttle in photosystem I. In those diatoms with both clade I and clade II flavodoxins, only clade II is induced under iron limitation. In those diatoms that lack clade II flavodoxin, the clade I flavodoxin is also induced by iron limitation, although at two orders of magnitude lower transcript abundances (Fig. 2B-C, Table S1). This suggests that in those diatoms with only clade-I flavodoxins, Fd replacement as a means of reducing iron requirements may play a less important role in adaptation to low iron conditions.

The knock-out of clade I flavodoxin indicates that the clade I flavodoxin partially mitigates oxidative stress under iron replete conditions (Fig. 3C). Further support for the role of clade I flavodoxin during oxidative stress comes from our observation that the clade I flavodoxins are distinguished from the clade II flavodoxins by replacement of cysteine with alanine at residue 103 (Fig. S1F). Cysteines are susceptible to oxidation under oxidative stress, which would be expected to inactivate the clade II protein.

The observation that the clade I gene is transcribed at orders of magnitude lower levels than the clade II gene suggests either that different transcriptional controls may regulate clade I flavodoxin transcript accumulation within a cell, or that oxidative stress may deplete a smaller fraction of the Fd pool compared to iron deprivation. Alternatively, low clade I transcript abundances in our bulk measurements may reflect the previously observed heterogeneity in the response of individual cells to H_2_O_2_ exposure (Mizrachi et al., 2020), our means of eliciting oxidative stress. Identification of potential selective pressures for retention of one or both flavodoxin clades will benefit from mechanistic studies with the two proteins to determine whether, as expected, they display different efficiencies under conditions that elicit susceptibility to oxidative stress, and whether their protein levels are differently controlled.

Our metatranscriptomes studies of eukaryotic phytoplankton communities identified both general patterns in transcriptional profiles of the clade I and II flavodoxin genes and highlighted specific responses of different diatoms. Clade II flavodoxins in diatoms in the wild are transcribed in environments where nitrate assimilation is decreased due to iron limitation, defined here by elevated N/Fe (>4 μm N / nm Fe). The different detected diatom genera displayed specific responses to conditions encountered along the two transects. The environmental *Chaetoceros* transcribed clade II flavodoxin only at putative iron-limiting stations, despite detection of this genus at additional iron-replete stations. In contrast, two other environmental diatom genera, *Pseudo-nitzschia* and *Fragilariopsis* were both numerically abundant based on their total transcript abundances (Fig. S5C), and yet the clade II transcript abundance for these two species remained relatively low throughout both transects, suggesting these species were not experiencing iron limitation at the time of sampling. Similarly, the pelagophyte *Pelagomonas* was also abundant across the transects and yet also transcribed clade II at relatively low levels (Fig. S5B-C). Importantly, several of the detected *Chaetoceros* species encode clade I flavodoxins (Fig. S4), which were either detected at relatively low levels or not detected at all along the transects indicating the specificity of transcription of the two flavodoxin clades in response to environmental conditions.

The NPSG is an iron-replete, high-light environment, conditions which enhance oxidative stress and may thus compromise Fd functionality. Consistent with this premise, the greatest relative abundance of clade I transcripts was detected within the gyre, with a majority of the transcripts assigned to *Fistulifera solaris*, which displayed a diel peak in transcript abundance each dawn (Fig. 5H). This rhythmic pattern was detected only with the clade I flavodoxin from *Fistulifera* suggesting either a genus-specific adaptation to high light or other oxidative stress generating processes, such as biotic interactions. Interestingly, genome-wide transcriptomic profiles of two model diatoms over the diel cycle, also suggest a diel rhythm to clade I flavodoxins transcript abundance. *Phaeodactylum tricornutum* clade I flavodoxin (Phatr3_J13706) expression peaked just before and after dusk (Smith et al., 2016), and the benthic diatom *Seminavis robusta* induced the clade I flavodoxins (sro1985_g309400, sro668_g166050) before dusk (Bilcke et al., 2021) (Table S1). The diel rhythmicity of clade I flavodoxins in both laboratory studies with model diatoms, as well as the diel rhythmicity in natural diatom communities emphasizes the potential circadian regulation of this protein in anticipation of the dawn, and photosynthetic-related oxidative stress.

Here we show that not all flavodoxins are responsive to low iron concentrations, and that a subset of flavodoxins (clade I flavodoxins) may be important in oxidative stress response. This specialization might enable diatoms and other stramenopiles to survive in stressful rapidly changing environments. The genetic duplication and functional differentiation of flavodoxin described here add a molecular mechanism that may facilitate diatom survival and adaptation to two major stresses – iron limitation and oxidative stress

## Material and Methods

### HMM search and phylogenetic tree construction

A custom-made flavodoxin hidden Markov model (hmm)-profile was generated (hmm-build; Eddy, 1998) from an alignment containing the Flavodoxin_1 seed alignment (Pfam id PF00258; pfam.xfam.org; Finn et al., 2016) amended with the flavodoxin amino acid sequences of *P. tricornutum*, *T. pseudonana* and *T. oceanica*e, using Multiple Alignment using Fast Fourier Transform (MAFFT) version 7.313 (Katoh et al., 2002; parameters: --localpair --maxiterate 100 –reorder; supplemental fasta file 1). Sequences from publicly available genomes and transcriptomes of over 500 marine protists, bacteria, archaea, and viruses (Coesel et al., 2021), were searched using the custom hmm-profile (e < 0.001; hmmsearch; Eddy, 1998), and hits were clustered at 99% identity by usearch (Edgar, 2010). For phylogenetic analysis, sequences were aligned with MAFFT (parameters: -- maxiterate 100 --reorder --leavegappyregion). A maximum-likelihood phylogenetic tree was built using RAxML version 8.2.4 (Stamatakis, 2014; parameters: -f a -m PROTGAMMAWAG - p 451325 -x 12345 -# 100; Fig. S1A-B, Fig. 1A). As previously noted (Caputi et al., 2019), the Pfam and HMM-search do not discriminate those sequences involved in photosynthetic metabolism from other homologous sequences. The phylogenetic tree was used to distinguish clades containing known photosynthetic flavodoxins from the outgroup clade containing nonphotosynthetic flavodoxin sequences (labeled in black and red respectively, Fig. S1A).

A stramenopile-focused phylogenetic analysis was generated by a similar search of 56 additional stramenopile sequences obtained from diverse data sources (Table S2). The sequences were identified using our custom flavodoxin hmm profile (e < 0.001; hmmsearch) and added to the original alignment using MAFFT (parameters: --add --localpair --maxiterate 100 --reorder -- leavegappyregion). A fast tree was generated by RAxML (parameters: -f E -p 271321 -m PROTGAMMAWAG) and used to prune branches from the “outgroup” area (left of the dashed line in Fig. S1A), removing most of the similar clades, and leaving some representatives to attract the environmental reads in the RAxML EPA analysis, described below. The sequences from *Tiarina fusus* were also removed from the tree, as this ciliate was reported to be heavily contaminated with prey (Lasek-Nesselquist and Johnson, 2019). Additional rogue sequences were detected by RogueNaRok by majority-rule consensus (MR; Aberer et al., 2013), and a ‘rawImprovement’ cut-off of > 0.85 was applied to sequences located within the four photosynthetic flavodoxin clades, whereas a more strict cutoff (> 0.2) was applied to the outgroup. The remaining 611 sequences were used to generate the stramenopile photosynthetic flavodoxin-focused tree (parameters: -f a -x 25114 -p 269321 -# 100 -m PROTGAMMAWAG) (Fig. S1C, Fig. 1B). All tree visualizations were performed in the Interactive Tree of Life version 5 (https://itol.embl.de/; Letunic and Bork, 2021).

### Culture growth conditions

All diatom isolates were obtained from the Provasoli-Guillard National Center for Culture of Marine Phytoplankton NCMA). Cultures were grown in 16:8 hours light:dark cycles and a light intensity of about 100 μmol photons·m^-2^·sec^-1^. *T. pseudonana* (CCMP1335), *T. oceanica* (CCMP1005), *Amphora coffeaeformis* (CCMP1405) were grown at 20 °C. *Chaetoceros sp*. (CCMP199), and *Cylindrotheca closterium* (CCMP340) were grown at 24 °C. All cultures were adapted to artificial sea water (either EASW or Aquil media) for at least a month before the experiments started. See Table S3 for each species media type and exact growth conditions. The following media were used: f/2 (Guillard and Ryther, 1962), L1 (Guillard and Hargraves, 1993), EASW (Berges et al., 2001), Aquil (Price et al., 1989). For iron limitation experiments, exponentially growing cells were centrifuged at 4000 rpm (5-10 min, 20-22 °C as specified in Table S3). Cells were washed three times with growth media lacking added Fe, and then diluted into either replete media, or media with no added Fe and 1.5 μM desferrioxamine B (DFB; iron chelator, Sigma). Cells were kept in semi-exponential state by dilutions in fresh media. The cultures were grown in polystyrene flasks to minimize iron contamination. For oxidative stress experiments, we determined the lethal dose for each species in preliminary small-scale experiments and used the lowest lethal concentration (250 μM for *Amphora*, and 200 μM for the other cultures). In contrast to the experiments used for transcriptome analysis, all experiments with *T. pseudonana* WT versus the KO lines were done in filtered sea water (FSW) from Puget Sound, supplemented with f/2, iron limitation was achieved by diluting exponentially growing cells into FSW supplemented with f/2 or f/2 without added iron.

### Cell counts and cell death

Samples of 150 μl from triplicate flasks were taken and measured immediately using Guava easyCyte 11HT Benchtop Flow Cytometer, excitation by 488 nm laser. Cells were detected by chlorophyl autofluorescence (680±30 nm) and forward scatter. Cell death was determined by positive Sytox Green (Invitrogen) staining, used at a final concentration of 1 μM. Samples were incubated in the dark for 30 min prior to measurement. Positive gating (525±30 nm) was based according to untreated stained cells and unstained cells.

### Photosynthetic efficiency

Photochemical yield of photosystem II (Fv/Fm) was determined with a Phyto-PAM fluorometer (Heinz Walz GmbH, Effeltrich, Germany) using 15 min dark-adapted cells. Triplicate samples were measured, 1 ml in 1 cm cuvette, each sample was measured 3 times with 30 sec intervals.

### RNA extraction, and transcriptome analysis

Cells were harvested by filtration onto 0.22 μm filters. Full details of the number of cells harvested per treatments, per species, and samples that fail library preparation are indicated in Table S3. Filters were snap-frozen in liquid nitrogen and stored at −80 °C until extraction. Total RNA was extracted from 0.22 μm membrane filters with the Zymo Direct-zol RNA MiniPrep Plus kit. RNA was quantified using Qbit, and 1900 ng per sample were sent to the Northwest Genomics Center (University of Washington). Poly-A-selection, library preparation and sequencing were performed at the Northwest Genomics Center with Illumina NextSeq and standard protocols. Sequence reads were trimmed using Trimmomatic 0.39 (Bolger et al., 2014) run in paired-end mode with cut adaptor and other Illumina-specific sequences (ILLUMINACLIP) set to TruSeq3-PE.fa:2:30:10:1:true, Leading and Trailing thresholds of 25, a sliding window trimming approach (SLIDINGWINDOW) of 4:15, an average quality level (AVGQUAL) of 20, and a minimum length (MINLEN) of 60. Reads were mapped to the genome of *T. pseudonana* or *T. oceanica* using Hisat2-2.1.0. We calculated the number of *T. pseudonana* and *T. oceanica* reads that aligned to the gene models from resulting sequence alignment map (SAM) files with aligned sequences used in subsequent analyses and normalized using transcripts per million (TPM). *De-novo* transcriptome assemblies were generated from RNA sequences extracted from isolate cultures of *Amphora, Chaetoceros*,and *Cylindrotheca*. Each species’ quality-controlled RNA sequencing data was assembled using Trinity v2.12.0 using default settings and with sequences provided as un-merged paired end reads. The pre-assembly RNA sequencing reads were then mapped back to their respective assemblies as a quality control step, and at least 85% of the reads from all cultures successfully mapped onto the resulting assemblies. The assembly resulted in 53,401 contigs for *Amphora*,37,507 contigs for *Chaetoceros*, and 73,553 contigs for *Cylindrotheca*.

The edgeR pipeline (Chen et al., 2010) was used to identify the transcriptional changes in different conditions for all five species, and visualized in MDS plot (Fig. S2D). EdgeR was used to detect differential expression of log2 transformed transcript levels based on pairwise comparisons between the control samples and the treated samples. A generalized linear model (GLM) quasi-likelihood F-test (QLTF) was used to test for significant differential expression (p < 0.01 and false discovery rate (FDR) < 0.05).

Flavodoxins genes of *T. pseudonana* and *T. oceanica* were taken from the genomes (gene ids: THAPSDRAFT_28635, THAOC_31152, THAOC_19008, THAOC_16623). For *Amphora, Chaetoceros* and *Cylindrotheca* the flavodoxins sequences of the same species, coastal isolates were used in blast against each species assembly (tblastn, -evalue 0.001). The queries and detected flavodoxins are presented in supplementary fasta file 5. These flavodoxins are present in the flavodoxins phylogenetic tree (Fig. 1B, indicated with red stars, Table S2, indicated by relevant expression – this study).

### TpFlav Gene

The gene sequence and amino acid sequence of *T. pseudonana* flavodoxin were obtained from the JGI genome portal (THAPSDRAFT_28635, transcript ID: EED91575).

### gRNA design for flavodoxin knockout

We combined Golden-Gate cloning using two sgRNAs (Hopes et al., 2017, 2016), with bacterial conjugation that allows transient transformation and removal of the Cas9 after a deletion is verified (Karas et al., 2015). Two sgRNAs were designed to cut ~120 nucleotides, including an FMN binding site and conserved region (Fig. S3A). Selection of CRISPR/Cas9 targets and estimating on-target score: Twenty bp targets with an NGG PAM were identified and scored for on-target efficiency using the Broad Institute sgRNA design program (www.broadinstitute.org/rnai/public/analysis-tools/sgrna-design), which utilizes the (Doench et al., 2014) on-target scoring algorithm. The sgRNAs that were chosen had no predicted off-targets: The full 20 nt target sequences and their 3’ 12 nt seed sequences were subjected to a nucleotide BLAST search against the *T. pseudonana* genome. Resulting homologous sequences were checked for presence of an adjacent NGG PAM sequence at the 3’ end. The 8 nt sequence outside of the seed sequence was manually checked for complementarity to the target sequence. In order for a site to be considered a potential off-target, the seed sequence had to match, a PAM had to be present at the 3’ end of the sequence and a maximum of three mismatches between the target and sequences from the BLAST search were allowed outside of the seed sequence.

### Plasmid construction using Golden Gate cloning

Golden Gate cloning was carried out as previously described (Weber et al., 2011), using a design similar to (Graff van Creveld et al., 2021; Hopes et al., 2016). The level 1 (L1) plasmid that enable conjugation was a kind gift from Irina Grouneva, Mackinder lab, University of York, UK (Nam et al., 2022). Golden Gate reactions for L1 and level 2 (L2) assembly were carried out, using 40 fmol of each component was included in a 20 μl reaction with 10 units of BsaI or BpiI and 10 units of T4 DNA ligase in ligation buffer. The reaction was incubated at 37 °C for 5 h, 50 °C for 5 min and 80 °C for 10 min. Five μl of the reaction was transformed into 50 μl of NEB 5α chemically competent *Escherichia coli*. The nourseothricin resistance gene (NAT) was PCR-amplified from pICH47732:FCP:NAT (Addgene #85984) using primers 1 and 2 (Table S5), and cloned into a pCR8/GW/ TOPO vector (ThermoFisher). The FCP promoter, NAT and FCP terminator level 0 (L0) modules (Hopes et al., 2016) were assembled into L1 pICH47751. The sgRNA scaffold was amplified from pICH86966_AtU6p_sgRNA_NbPDS (Nekrasov et al., 2013) with sgRNA sequences integrated through the forward primers (3-5, Table S5). Together with the L0 U6 promoter (pCR8/GW:U6, Addgene #85981), the two sgRNAs were assembled into L1 destination vectors pICH47761 and pICH47772. Insertion areas in the L1 plasmids were sequenced (6-7, Table S5). L2 assembly: L1 modules and annealed oligonucleotides (8-9, Table S5) assembled into the L2 destination vector pAGM4723-Del (Addgene #112207). The final plasmids, L2_Conj_Cas_NAT_Flav were screened by colony PCR, and the sgRNAs area was sequenced (10-12, Table S5).

### *T. pseudonana* transformation by bacterial conjugation

The L2_Conj_Cas_NAT_Flav plasmid was delivered to *T. pseudonana* cells via conjugation from *E. coli* TOP10 cells as previously described (Karas et al., 2015). The mobilization helper plasmid pTA-Mob, containing all genes necessary for the conjugative transfer of oriT-containing plasmids, was a gift from Rahmi Lale (Strand et al., 2014), and was transformed into the cells. *E. coli* pTA-Mob electroporation-competent cells (50 μl) were transformed with 100 ng of the L2_Conj_Cas_NAT_Flav plasmid and used for conjugative delivery of the CRISPR/Cas9 plasmid to the diatom cells. Overnight *E.coli* cultures were inoculated into 50 ml LB + kanamycin (50 mg l^-1^) and gentamicin (20 mg l^-1^), and grown with shaking at 37 °C to an OD_600_ of about 0.3. About 40 ml were harvested by 10 min centrifugation at 4000 rpm, 10 °C, and resuspended in 400 μl SOC. *T. pseudonana* cells at about 9·10^5^ cells/ml and Fv/Fm of 0.57 were harvested by centrifugation (4000 rpm, 10 min, 18 °C). About 8·10^7^ cells were suspended in 500 μl of FSW. *T. pseudonana* and *E. coli* cells were mixed, and plated on two plates of 50% FSW L1, 0.8% (w/v), 5% LB (v/v). After drying, the plates were incubated for 90 min at 30 °C in the dark, then moved to the regular growth chamber (20 °C, about 100 μmol photons·m^-2^·sec^-1^) overnight. Then 1 ml of *T. pseudonana* medium was added to each plate and cells were scraped. Cells from each plate were plated on two plates of 50% FSW L1, 0.8% (w/v), nourseothricin 50 or 100 mg·ml^-1^, 0.8% agar plate (selection plates) and incubated at 18 °C. Colonies appeared after about two weeks.

### Selection of knockout lines

Colonies were scanned for the size of flavodoxin amplicon (primers 13-14, Table S5), colonies exhibiting double-bands, representing both WT (676 bp) and edited (~530 bp) flavodoxin (probably heterozygotes or mosaic colonies), were re-streaked onto fresh solid medium containing 100 μg·ml^-1^ nourseothricin. Daughter colonies were scanned for the size of flavodoxin amplicon, colonies exhibiting a single band representing bi-allelic edited flavodoxin were selected for further use. Several colonies exhibiting a single band, either indicative of WT or edited flavodoxins, were transferred into non-selective media for few weeks. Cells were spread onto non-selective solid-media, single colonies were picked and tested for antibiotic resistance. The flavodoxin gene of nonresistant colonies was sequenced to locate the exact deletion for KO or verify no DNA editing as a control (primers 14-15 Table S5). WT and two colonies without deletion in the flavodoxin (colonies 5 and 16) are referred here as “WT”. Three colonies with deletion in the flavodoxin gene (9, 14, 1) are named here “KO”.

### Specific cruises sample collection

#### Diel 1

Samples were collected during 25 July to 5 August 2015 aboard the R/V *Kilo Moana*, cruise KM1513 from ~156.5°W, 24.5° N. About 7 L seawater samples were pre-filtered through 100 μm nylon mesh and collected onto 142 mm 0.2 μm polycarbonate filters using a peristaltic pump. The cruise was previously described (Wilson et al., 2017), additional cruise information and associated data can be found online: https://simonscmap.com/catalog/cruises/KM1513.

#### Gradients 1

Samples were collected during 19 April to 3 May 2016 aboard the R/V *Ka’imikai-O-Kanaloa*, cruise KOK1606, from ~23.5 - 37 °N, 158 °W (Juranek et al., 2020). About 6-10 L seawater samples were pre-filtered through a 200 μm nylon mesh and collected by sequential filtration through a 3 μm and a 0.2 μm polycarbonate filters using a peristaltic pump. In this study, we combined the two size-fraction in the analysis. Additional cruise information and associated data can be found online: https://simonscmap.com/catalog/cruises/KOK1606.

#### Gradients 2

Samples were collected during 27 May to 13 June 2017 aboard the R/V *Marcus G. Langseth*, cruise MGL1704, from ~26 - 41 °N, 158 °W (Juranek et al., 2020). About 6-10 L seawater samples were pre-filtered through a 100 μm nylon mesh and collected as above. Additional cruise information and associated data can be found online: https://simonscmap.com/catalog/cruises/MGL1704.

#### Gradients 2 on-deck incubations

At 37 °N, 158 °W a total of 20 L seawater was collected into replicate polycarbonate carboys from 15 m depth and incubated at *in situ* temperature for 96 h in on-deck, temperature-controlled incubators screened with 1/8 inch light blue acrylic panels to approximate *in situ* light levels at 15 m. Triplicate carboys were sampled immediately (T = 0) or amended with of 1 nM FeCl_3_ (+Fe), of 5 μM NO_3_ and 0.5 μM PO_4_ (+N +P). Triplicate carboys with no amendment served as a control. After 4 days, the samples were filtered in the same way as the transect samples (Lambert et al., 2022).

All metatranscriptomic samples were flash frozen in liquid nitrogen and subsequently stored at −80 °C until further processing. RNA extraction and sequencing was previously described (Durham et al., 2019; Lambert et al., 2022). Briefly, RNA was extracted with a set of 14 internal mRNA standards in the extraction buffer. After DNase treatment, purification and quantification, the eukaryotic mRNAs were poly(A)-selected, sheared and used to construct complementary DNA libraries. Sequences were *do novo* assembled and functionally and taxonomically annotated. The metatranscriptome data from the Diel1 are available through the NCBI’s SRA under BioProject PRJNA492142, Gradients 1 under BioProject PRJNA690573, and Gradients 2 under BioProject PRJNA690575.

### Environmental metadata sourcing and processing

Dissolved iron and nitrogen concentrations were measured during the Gradients cruises (Gradoville et al., 2020; Juranek et al., 2020; Park et al., 2022; Pinedo-González et al., 2020) and retrieved from the Simons Collaborative Marine Atlas Project pycmap API (CMAP; https://simonscmap.com/).

### Flavodoxins in the metatranscriptomes

The flavodoxin detection and quantification bioinformatic workflow is shown in Fig. S5A. For the three cruises, environmental metatranscriptome contigs with homology to flavodoxin were recruited to the custom-made hmm-profile described above (e < 0.001; hmmsearch). The environmental sequences were aligned to the flavodoxin alignment (stramenopile-enhanced, supplementary fasta file 3) using MAFFT (parameters: --add --localpair --maxiterate 100 --reorder), and placed on the flavodoxin reference phylogenetic tree (Fig. 1B, Fig. S1C) with the additional stramenopiles by RAxML Evolutionary Placement Algorithm (EPA; Barbera, 2016; parameters: -f v -m PROTGAMMAWAG). Environmental reads with like_weight_ratio > 0.8 that were placed on clade I or II stramenopiles at genus level or more specific were kept (Fig. S4). The clade and genus were determined from the flavodoxin reference phylogenetic tree. Reads per liter from each sample were summed to the clade and genus level, and divided by all the reads of the taxonomic order of the contigs, as determined by Lowest Common Ancestor (LCA) using the LCA algorithm in DIAMOND in conjunction with NCBI taxonomy (Buchfink et al., 2014), as was previously described (Groussman et al., 2021). The normalized counts were averaged for the replicates, and the two size-fractions were summed. Short reads from the incubation experiments were mapped against the flavodoxin contigs from Gradients 2 using kallisto (Bray et al., 2016) and normalized to reads per liter using synthetic standards, as in (Coesel et al., 2021; Durham et al., 2019). The normalized reads were aggregated and summed to genus level, the replicates were averaged and the two size-fractions were summed. In Fig. 5G the reads are normalized per clade and genus. The difference between this method and the underway stations, is shown in Fig. S5A. **Determining significant diel periodicity** : Significant periodicity of the Diel 1 contigs was taken from Groussman et al., 2021, with the Rhythmicity Analysis Incorporating Non-parametric Methods (RAIN) package implemented in R (Thaben and Westermark, 2014). The p-values from RAIN analyses were ranked and corrected at an FDR < 0.05 using the Benjamini-Hochberg false-discovery rate method.

## Supporting information

Supplemental Fasta files

## Data availability

All relevant data supporting the findings of the study are available in this article and its Supplementary files. All the new RNAseq data will be publicly available on NCBI.

## Acknowledgments

We thank the crew and scientific party of the *R/V Kilo Moana, R/V Kaimikai O Kanaloa* and *R/V Langseth*, and the operational staff of the Simons Collaboration on Ocean Processes and Ecology (SCOPE) program for logistical support. We thank Bryn Durham for assisting with on-deck incubation, Aidan DeHan for assistance in physiological measurements, and Zinka Bartolek for fruitful discussions. This work was supported by a grant from the Simons Foundation (SCOPE Award ID 721244 to E. V. A.).

## Supplementary tables

**Table S1.** Diatoms flavodoxins expression from the literature. Species name, original paper, and the specific information about flavodoxin gene ID and the expression in each paper are indicated.

**Table S2.** Information about the species and strains with clade I or clade II flavodoxins. Includes sequence id, taxonomy, data type (genome or transcriptome), isolate location and growth temperature (from culture collections NCMA or RCC when possible, from manuscript describing the isolate when possible). Reference for relevant transcriptome when available, number of flavodoxins from each clade in the phylogenetic trees (# clade I / II). Number of similar flavodoxins that were remove from the alignment due to high sequence similarity (Removed* clade I / II). Strains presented in Fig. 1A (Yes), or within the stramenopiles added for Fig. 1B.

**Table S3.** Diatom species, with information about the exact isolates, growth and harvesting conditions for the different transcriptomes done in this study.

**Table S4.** Stramenopiles flavodoxins transcription in the North Pacific, data presented in Fig. 5. Stramenopiles flavodoxins transcription across the Gradients 1 transect, R/V Kaimikai O Kanaloa, KOK1606 (April/May 2016), and Gradients 2 transect, R/V Langseth, MGL1704. Flavodoxin sequence reads per liter were summed at the genus level and normalized to the cumulative number of reads per liter at the order level to compare between genera. Stramenopiles flavodoxin transcription following nutrient enrichment incubations conducted at Gradients 2, station 11, 37 °N. Water samples were sample at T=0, and incubated with no added nutrients (Control) or with 1 nM FeCl_3_ (+Fe), or 5 μM NO_3_ and 0.5 μM PO_4_ (+N +P) sampled for metatranscriptomes after 4 days. Transcripts per liter were summed at the genus level.

**Table S5.** Primers used for Golden Gate cloning, plasmid verification, and deletion scan and sequencing of *T. pseudonana* flavodoxin gene.

## Supplementary figure legends

**Figure S1.** Flavodoxins in publicly available databases

**A.** Maximum-likelihood (RAxML) tree of flavodoxins. Names of organisms with non-photosynthetic flavodoxins proteins or proteins with flavodoxin domains labeled in red and previously identified photosynthetic flavodoxins labeled in black. Photosynthetic flavodoxins (right side of dashed line) are presented in Fig. 1A. Thick branch lines represent bootstrap support greater than 0.7. Different clades are colored as in Fig. 1: clade I (orange), clade II (purple), bacteria/green algae clade (pink), cyanobacteria/ dinoflagellate/green algae clade (green).

**B.** Photosynthetic flavodoxins of the same maximum-likelihood (RAxML) phylogenetic tree presented in A. Thick branch lines represent bootstrap support greater than 0.7. Clades are marked with colored branches as in A. Colored strip represents taxonomy, color coded on the figure. Isolate habitat of stramenopiles is indicated: blue circles (open ocean), brown full rectangles (coastal), brown open rectangles (other – benthic, hypersaline, freshwater or tidal mud), see Table S2 for isolate location. Taxonomic labels are black for previously identified photosynthetic flavodoxins and gray for other species.

**C.** Unrooted maximum-likelihood (RAxML) phylogenetic tree of clade I and II flavodoxins after inclusion of 56 stramenopile strains (see table S2 for list of added strains). Thickness of the branch lines represent bootstrap support (see legend). Clades are marked with colored branches as in panels A and B. Colored strip represents taxonomy. Non-photosynthetic flavodoxins as well as flavodoxin domains in other proteins labeled in red, previously identified photosynthetic flavodoxins labeled in black. Red stars represent flavodoxins with expression experimentally tested under iron limitation. This is the full tree of Fig. 1B.

**D.** Number of flavodoxins from each clade detected in stramenopiles. Each row is a strain. Left column represent the taxonomy, white – stramenopiles that are not diatoms, diatoms: grays – pennates, dark raphid, light araphid, greens: centric, dark – radial, light – polar. The other columns represent the number of flavodoxins detected from each clade (center – clade I, right – clade II), white – 0, darkest green – 8.

**E.** Schematic phylogeny within the stramenopiles group, adapted from (Derelle et al., 2016), Pinguiophyceae placed according to (Kawachi et al., 2002). Groups with detected clade I flavodoxins are marked with orange asterisk, groups with only clade II flavodoxins detected are marked with black X, groups with no flavodoxin detected are marked with gray circle. The branches are colored accordingly. Notably, the different groups are not evenly represented in our dataset. There are 67 centric diatoms (13 with no clade I flavodoxin), while there are 3 Chrysophyceae and one *Dictyocha*.

**F.** Conserved FMN binding sites in stramenopiles flavodoxins. Top row: Conserved sequences surrounding the FMN molecule according to structure of a red alga *Chondrus crispus* flavodoxin (Fukuyama et al., 1992). Amino-acids are numbered according to *C. crispus*.Asterisks indicate the residues whose side chains flank the FMN, amino acids whose side-chains form hydrogen bond with the FMN are underlined, amino acids whose backbone form a hydrogen bond with the FMN are marked with gray rectangles. Middle, bottom rows: Sequence logos detailing alignment conservation among each clade of flavodoxins of diatoms, and other stramenopiles. Black arrows indicate amino-acids 57 and 103 discussed in the text. Sequence logos were created with the R package Ggseqlogo (Wagih, 2017), using alignments present in supplementary fasta 4.

**Figure S2.** Diatoms cultures transcriptomes

**A.** Experimental design and pictures of representative flasks at the time of harvesting for transcriptomes. The full experiment was done in biological triplicates. For each measurement, samples were taken from the triplicates.

**B.** Cell counts, measured by flow cytometry of the 5 diatom species during the experiment. Iron replete (control, gray circles), iron limitated (-Fe, green triangles), line breaks indicate culture dilution into the same growth media. A subset of replete cultures was treated with H_2_O_2_ ~ 1.5 h before harvest for transcriptomes (light blue squares).

**C.** Photosynthetic efficiency (Fv/Fm) of the same experiment. Colors and symbols as in B.

**D.** Multidimensional scaling plot (MDS) of the 5 diatoms transcriptomes, generated by EdgeR (Robinson et al., 2009). Colors and symbols as in B and C. Some samples were not sequenced due to RNA loss during library preparation: 1 control *Amphora*, 2 control, 1 iron limitation *Cylindrotheca*, 1 control, 1 iron limitation, 2 H_2_O_2_ treated *Chaetoceros*.

**Figure S3.** Flavodoxin knock-out in *T. pseudonana*

**A.** *T. pseudonana* flavodoxin (TpFlav) DNA and amino acid sequence, exons (teal), FMN binding sites (orange), amino acids with side chains that flank the FMN (*) and sgRNAs (red) are marked. Targeted deletion in gray background. FMN binding sites according to *C. crispus* flavodoxin (Fukuyama et al., 1992).

**B.** PCR of the deletion area of *TpFlav*. Lanes from left to right: DNA ladder (size in bp), WT, two WT-like clones (5, 16), three knock-out (Flav KO) clones (9, 14,1), negative control.

**C.** Sequence of targeted deletion in *TpFlav* in WT, WT-like (16, 5), Flav KO (1, 9, 14) cells, aligned to the DNA and amino acid sequence. Amino-acids with side chains that flank the FMN (*) and sgRNAs (red) are marked. The deletion is marked by red background. Alignments generated and visualized with the SnapGene software.

**D-H.** The three independent WT (grays) and Flav KO (oranges) sets of lines were grown under iron-replete (**D**, **E**, **H**) and iron-limiting (**F**, **G**) conditions for several days. **D**, **F.** Cell abundance, measured by flow cytometry. **E**, **G.** Photosynthetic efficiency, measured by phytoPAM. Individual measurements are marked in symbols, means of triplicates in lines. **H.** Percentage of Sytox Green-positive (dead) cells, measured by flow cytometry 24 h after treatment with 100 or 200 μM H_2_O_2_. Each box plot represents a cell line, single measurements are marked.

**Figure S4.** Flavodoxins in the North Pacific Ocean

Maximum-likelihood (RAxML) tree of flavodoxins, with placements of environmental reads from the North Pacific Ocean. This tree is also represented in Fig. 1B, Fig. S1C and Fig. 4B. Branches are orange for clade I, purple for clade II. Colored strip represents taxonomy, color coded on the figure. Black symbols represent diatom class: closed triangles – raphid pennates; open triangles – araphid pennates; closed circles – polar centrics; open circles-radial centrics. The genus and species names are in black for diatoms, or gray for all other species. Stars represent flavodoxins with expression experimentally tested, open stars indicate induction under iron limitation, red-filled stars indicate no induction under iron limitation. Environmental reads (contigs) placed confidently (like_weight_ratio > 0.8) at genus level or below are shown with pie charts color-coded by cruise. Pie size is proportional to total number of contigs.

**Figure S5.** Transcriptional patterns of flavodoxin genes in the North Pacific Ocean

**A.** Metatranscriptomes workflow chart for detection annotation and quantification of environmental flavodoxins. Left gray box represents analysis prior to this study, RNA extraction, sequencing, assembly, annotation, and quantification (Coesel et al., 2021; Durham et al., 2019; Groussman et al., 2021; Lambert et al., 2022). Central part represents the specific search, annotation and quantification of flavodoxins. Green arrow represents steps common to analysis of metatranscriptomes from diel 1, Gradients 1 and 2, the datasets used to quantify flavodoxin reads are highlighted in yellow boxes. Right orange box represents the process specific to the Gradients 2 on-deck incubations.

**B.** Heatmaps of relative transcript abundances of non-diatom stramenopile flavodoxins across the Gradients 1, and Gradients 2 transects, on-deck incubation experiment and Diel 1 cruise. Flavodoxin sequence reads per liter were summed at the genus level and normalized to the cumulative number of reads per liter at the order level to compare between genera. Note the different color-scale ranges. In the incubation experiment transcripts per liter were summed at the genus level and normalized to cumulative number of reads per liter at the genus level to compare within each genus. Upper bar represents daytime (yellow) or night (gray). Side bars represent flavodoxin clade I (orange) or clade II (purple). Each panel was color-scaled separately for better visualization (see separate color keys). White rows indicate genera-level flavodoxins that were not detected in a given cruise.

**C.** Cumulative counts per genus across the Gradients transects and the Diel 1 cruise. Stramenopiles genera with detected flavodoxins are presented. Each cruise and diatom or non-diatoms are color-scaled separately for visualization.

## Supplemental fasta files

1. Amino-acid sequence alignments of flavodoxin PF00258 and *P. tricornutum T. pseudonana T. oceanica*, used to generate the hmm-profile.
2. Amino-acid sequence alignments of flavodoxins used to create the phylogenetic tree in Fig. 1A.
3. Amino-acid sequence alignments of flavodoxins used to create the phylogenetic tree in Fig. 1B.
4. Amino-acid sequence alignments of stramenopiles flavodoxins aligned to *Chondrus crispus* flavodoxin, used for Fig. S1F. All the above are aligned and trimmed.
5. Flavodoxins sequences used to detect flavodoxins in *Amphora, Cylindrotheca* and *Chaetoceros* transcriptomes, and detected flavodoxins in those species.

## References

Aberer AJ, Krompass D, Stamatakis A. 2013. Pruning rogue taxa improves phylogenetic accuracy: An efficient algorithm and webservice. Syst Biol 62:162–166. doi: 10.1093/sysbio/sys078

Ashkezari MD, Hagen NR, Denholtz M, Neang A, Burns TC, Morales RL, Lee CP, Hill CN. 2021. Simons Collaborative Marine Atlas Project (Simons CMAP): an open-source portal to share, visualize and analyze ocean data. Limnol Oceanogr Methods 1–14. doi:10.1002/lom3.10439

Barbera P. 2016. Efficient and Massively Parallel Implementation of the Evolutionary Placement Algorithm. University of the State of Baden-Wuerttemberg and National Research Center of the Helmholtz Association.

Barbera P, Kozlov AM, Czech L, Morel B, Darriba D, Flouri T, Stamatakis A. 2019. EPA-ng: Massively Parallel Evolutionary Placement of Genetic Sequences. Syst Biol 68:365–369. doi:10.1093/sysbio/syy054

Behrenfeld MJ, Milligan AJ. 2013. Photophysiological expressions of iron stress in phytoplankton. Ann Rev Mar Sci 5:217–246. doi:10.1146/annurev-marine-121211-172356

Berges JA, Franklin DJ, Harrison PJ. 2001. Evolution of an artificial seawater medium: Improvements in enriched seawater, artificial water over the last two decades. J Phycol 37:1138–1145. doi:10.1046/j.1529-8817.2001.01052.x

Bilcke G, Osuna-Cruz CM, Silva MS, Poulsen N, D’hondt S, Bulankova P, Vyverman W, de Veylder L, Vandepoele K. 2021. Diurnal transcript profiling of the diatom Seminavis robusta reveals adaptations to a benthic lifestyle. Plant J. doi:10.1111/tpj.15291

Blanco NE, Ceccoli RD, Segretin ME, Poli HO, Voss I, Melzer M, Bravo-Almonacid FF, Scheibe R, Hajirezaei MR, Carrillo N. 2011. Cyanobacterial flavodoxin complements ferredoxin deficiency in knocked-down transgenic tobacco plants. Plant J 65:922–935. doi:10.1111/j.1365-313X.2010.04479.x

Boyd P, LaRoche J, Gall M, Frew R, McKay RML. 1999. Role of iron, light, and silicate in controlling algal biomass in subantarctic waters SE of New Zealand. J Geophys Res Ocean 104:13395–13408. doi:10.1029/1999jc900009

Boyd PW, Jickells T, Law CS, Blain S, Boyle EA, Buesseler KO, Coale KH, Cullen JJ, de Baar HJW, Follows M, Harvey M, Lancelot C, Levasseur M, Owens NPJ, Pollard R, Rivkin RB, Sarmiento J, Schoemann V, Smetacek V, Takeda S, Tsuda A, Turner S, Watson AJ. 2007. Mesoscale iron enrichment experiments 1993-2005: synthesis and future directions. Science 315:612–617. doi:10.1126/science.1131669

Bray NL, Pimentel H, Melsted P, Pachter L. 2016. Near-optimal probabilistic RNA-seq quantification. Nat Biotechnol 34:525–527. doi:10.1038/nbt.3519

Buchfink B, Xie C, Huson DH. 2014. Fast and sensitive protein alignment using DIAMOND. Nat Methods 12:59–60. doi:10.1038/nmeth.3176

Cammack R. 1982. Evolution and Diversity in the Iron-Sulphur Proteins. Chem Scr.

Campbell IJ, Bennett GN, Silberg JJ. 2019. Evolutionary Relationships Between Low Potential Ferredoxin and Flavodoxin Electron Carriers. Front Energy Res 7:1–18. doi:10.3389/fenrg.2019.00079

Caputi L, Carradec Q, Eveillard D, Kirilovsky A, Pelletier E, Pierella Karlusich JJ, Rocha Jimenez Vieira F, Villar E, Chaffron S, Malviya S, Scalco E, Acinas SG, Alberti A, Aury J, Benoiston A, Bertrand A, Biard T, Bittner L, Boccara M, Brum JR, Brunet C, Busseni G, Carratalà A, Claustre H, Coelho LP, Colin S, D’Aniello S, Da Silva C, Del Core M, Doré H, Gasparini S, Kokoszka F, Jamet J, Lejeusne C, Lepoivre C, Lescot M, Lima-Mendez G, Lombard F, Lukeš J, Maillet N, Madoui M, Martinez E, Mazzocchi MG, Néou MB, Paz-Yepes J, Poulain J, Ramondenc S, Romagnan J, Roux S, Salvagio Manta D, Sanges R, Speich S, Sprovieri M, Sunagawa S, Taillandier V, Tanaka A, Tirichine L, Trottier C, Uitz J, Veluchamy A, Veselá J, Vincent F, Yau S, Kandels-Lewis S, Searson S, Dimier C, Picheral M, Bork P, Boss E, Vargas C, Follows MJ, Grimsley N, Guidi L, Hingamp P, Karsenti E, Sordino P, Stemmann L, Sullivan MB, Tagliabue A, Zingone A, Garczarek L, D’Ortenzio F, Testor P, Not F, D’Alcalà MR, Wincker P, Bowler C, Iudicone D, Acinas SG, Bork P, Boss E, Bowler C, Vargas C, Follows MJ, Gorsky G, Grimsley N, Hingamp P, Iudicone D, Jaillon O, Kandels-Lewis S, Karp-Boss L, Karsenti E, Krzic U, Not F, Ogata H, Pesant S, Raes J, Reynaud EG, Sardet C, Sieracki M, Speich S, Stemmann L, Sullivan MB, Sunagawa S, Velayoudon D, Weissenbach J, Wincker P. 2019. Community-Level Responses to Iron Availability in Open Ocean Plankton Ecosystems. Global Biogeochem Cycles 33:391–419. doi: 10.1029/2018gb006022

Carradec Q, Pelletier E, Da Silva C, Alberti A, Seeleuthner Y, Blanc-Mathieu R, Lima-Mendez G, Rocha F, Tirichine L, Labadie K, Kirilovsky A, Bertrand A, Engelen S, Madoui M-A, Yoshikawa G, Romac S, Richter DJ, Poulain J, Dimier C, Kandels-Lewis S, Picheral M, Searson S, Jaillon O, Aury J-M, Karsenti E, Sullivan MB, Sunagawa S, Bork P, Not F, Hingamp P, Raes J, Guidi L, Ogata H, de Vargas C, Iudicone D, Bowler C, Meheust R, Wincker P. 2018. A global oceans atlas of eukaryotic genes. Nat Commun 9. doi:10.1038/s41467-017-02342-1

Ceccoli RD, Blanco NE, Medina M, Carrillo N. 2011. Stress response of transgenic tobacco plants expressing a cyanobacterial ferredoxin in chloroplasts. Plant Mol Biol 76:535–44. doi:10.1007/s11103-011-9786-9

Chen Y, McCarthy D, Ritchie M, Robinson M, Smyth G. 2010. edgeR: differential expression analysis of digital gene expression data. BioconductorFhcrcOrg.

Coesel SN, Durham BP, Groussman RD, Hu SK, Caron DA. 2021. Diel transcriptional oscillations of light-sensitive regulatory elements in open-ocean eukaryotic plankton communities. Proc Natl Acad Sci 118:1–12. doi: 10.1073/pnas.2011038118/-/DCSupplemental.Published

Derelle R, López-García P, Timpano H, Moreira D. 2016. A Phylogenomic Framework to Study the Diversity and Evolution of Stramenopiles (=Heterokonts). Mol Biol Evol 33:2890–2898. doi:10.1093/molbev/msw168

DiTullio GR, Geesey ME, Mancher JM, Alm MB, Riseman SF, Bruland KW. 2005. Influence of iron on algal community composition and physiological status in the Peru upwelling system. Limnol Oceanogr 50:1887–1907. doi:10.4319/lo.2005.50.6.1887

Doench JG, Hartenian E, Graham DB, Tothova Z, Hegde M, Smith I, Sullender M, Ebert BL, Xavier RJ, Root DE. 2014. Rational design of highly active sgRNAs for CRISPR-Cas9-mediated gene inactivation. Nat Biotechnol 32:1262–1267. doi:10.1038/nbt.3026

Dore JE, Letelier RM, Church MJ, Lukas R, Karl DM. 2008. Summer phytoplankton blooms in the oligotrophic North Pacific Subtropical Gyre: Historical perspective and recent observations. Prog Oceanogr 76:2–38. doi:10.1016/j.pocean.2007.10.002

Durham BP, Boysen AK, Carlson LT, Groussman RD, Heal KR, Cain KR, Morales RL, Coesel SN, Morris RM, Ingalls AE, Armbrust EV. 2019. Sulfonate-based networks between eukaryotic phytoplankton and heterotrophic bacteria in the surface ocean. Nat Microbiol 4:1706–1715. doi:10.1038/s41564-019-0507-5

Eddy SR. 1998. Profile hidden Markov models. Bioinformatics 14:755–763. doi:10.1093/bioinformatics/14.9.755

Edgar RC. 2010. Search and clustering orders of magnitude faster than BLAST. Bioinformatics 26:2460–2461. doi: 10.1093/bioinformatics/btq461

Erdner DL. 1997. Characterization of ferredoxin and flavodoxin as molecular indicators of iron limitation in marine eukaryotic phytoplankton. Massachusetts Institute of Technology, Woods Hole Oceanographis Institue.

Erdner DL, Anderson DM. 1999. Ferredoxin and flavodoxin as biochemical indicators of iron limitation during open-ocean iron enrichment. Limnol Oceanogr 44:1609–1615. doi:10.4319/lo.1999.44.7.1609

Finn RD, Bateman A, Clements J, Coggill P, Eberhardt RY, Eddy SR, Heger A, Hetherington K, Holm L, Mistry J, Sonnhammer ELL, Tate J, Punta M. 2014. Pfam: The protein families database. Nucleic Acids Res 42:222–230. doi: 10.1093/nar/gkt1223

Finn RD, Coggill P, Eberhardt RY, Eddy SR, Mistry J, Mitchell AL, Potter SC, Punta M, Qureshi M, Sangrador-Vegas A, Salazar GA, Tate J, Bateman A. 2016. The Pfam protein families database: Towards a more sustainable future. Nucleic Acids Res 44:D279–D285. doi:10.1093/nar/gkv1344

Fukuyama K, Matsubara H, Rogers LJ. 1992. Crystal structure of oxidized flavodoxin from a red alga Chondrus crispus refined at 1.8 Å resolution. Description of the flavin mononucleotide binding site. J Mol Biol 225:775–789. doi:10.1016/0022-2836(92)90400-E

Gradoville MR, Farnelid H, White AE, Turk-Kubo KA, Stewart B, Ribalet F, Ferrón S, Pinedo-Gonzalez P, Armbrust EV, Karl DM, John S, Zehr JP. 2020. Latitudinal constraints on the abundance and activity of the cyanobacterium UCYN-A and other marine diazotrophs in the North Pacific. Limnol Oceanogr 65:1858–1875. doi:10.1002/lno.11423

Graff van Creveld S, Ben-Dor S, Mizrachi A, Alcolombri U, Hopes A, Mock T, Rosenwasser S, Vardi A. 2021. Biochemical Characterization of a Novel Redox-Regulated Metacaspase in a Marine Diatom. Front Microbiol 12. doi:doi: 10.3389/fmicb.2021.688199Biochemical

Graff van Creveld S, Rosenwasser S, Levin Y, Vardi A. 2016. Chronic iron limitation confers transient resistance to oxidative stress in marine diatoms. Plant Physiol 172:968–979. doi:10.1104/pp.16.00840

Groussman RD, Coesel SN, Durham BP, Armbrust EV. 2021. Diel-Regulated Transcriptional Cascades of Microbial Eukaryotes in the North Pacific Subtropical Gyre. Front Microbiol 12:1–15. doi:10.3389/fmicb.2021.682651

Groussman RD, Parker MS, Armbrust EV. 2015. Diversity and evolutionary history of iron metabolism genes in diatoms. PLoS One 10:e0129081. doi:10.1371/journal.pone.0129081

Gueneau P, Morel F, Laroche J, Erdner D. 1998. The petF region of the chloroplast genome from the diatom Thalassiosira weissflogii: Sequence, organization and phylogeny. Eur J Phycol 33:203–211. doi:10.1080/09670269810001736703

Guillard R, Ryther J. 1962. Studies of marine planktonic diatoms. I. Cyclotella nana Hustedt, and Detonula confervacea (cleve) Gran. Can J Microbiol 8:229–239.

Guillard RRL, Hargraves PE. 1993. Stichochrysis immobilis is a diatom, not a chrysophyte. Phycologia 32:234–236. doi:10.2216/i0031-8884-32-3-234.1

Hehenberger E, Burki F, Kolisko M, Keeling PJ. 2016. Functional Relationship between a Dinoflagellate Host and Its Diatom Endosymbiont. Mol Biol Evol 33:2376–2390. doi: 10.1093/molbev/msw109

Hopes A, Nekrasov V, Belshaw N, Grouneva I, Kamoun S, Mock T. 2017. Genome editing in diatoms using CRISPR-Cas to induce precise bi-allelic deletions. bio-protocole 7:23. doi:10.21769/bioprotoc.2625

Hopes A, Nekrasov V, Kamoun S, Mock T. 2016. Editing of the urease gene by CRISPR-Cas in the diatom Thalassiosira pseudonana. Plant Methods 12:49. doi:10.1186/s13007-016-0148-0

Imanian B, Keeling PJ. 2007. The dinoflagellates Durinskia baltica and Kryptoperidinium foliaceum retain functionally overlapping mitochondria from two evolutionarily distinct lineages. BMC Evol Biol 7:1–11. doi:10.1186/1471-2148-7-172

Imanian B, Pombert JF, Keeling PJ. 2010. The complete plastid genomes of the two “Dinotoms” Durinskia baltica and Kryptoperidinium foliaceum. PLoS One 5. doi:10.1371/journal.pone.0010711

Imlay JA. 2006. Iron-sulphur clusters and the problem with oxygen. Mol Microbiol 59:1073–1082. doi:10.1111/j.1365-2958.2006.05028.x

Jeanjean R, Zuther E, Yeremenko N, Havaux M, Matthijs HCP, Hagemann M. 2003. A photosystem 1 psaFJ-null mutant of the cyanobacterium Synechocystis PCC 6803 expresses the isiAB operon under iron replete conditions. FEBS Lett 549:52–56. doi: 10.1016/S0014-5793(03)00769-5

Jones KL. 1988. Analysis of Ferredoxin and Flavodoxin in Anabaena and Trichodesmium Using Fast Protein Liquid Chromatography. Portland State University.

Juranek LW, White AE, Dugenne M, Henderikx Freitas F, Dutkiewicz S, Ribalet F, Ferrón S, Armbrust EV, Karl DM. 2020. The Importance of the Phytoplankton “Middle Class” to Ocean Net Community Production. Global Biogeochem Cycles 34. doi:10.1029/2020GB006702

Karas BJ, Diner RE, Lefebvre SC, McQuaid J, Phillips APR, Noddings CM, Brunson JK, Valas RE, Deerinck TJ, Jablanovic J, Gillard JTF, Beeri K, Ellisman MH, Glass JI, Hutchison III C a., Smith HO, Venter JC, Allen AE, Dupont CL, Weyman PD. 2015. Designer diatom episomes delivered by bacterial conjugation. Nat Commun 6:6925. doi:10.1038/ncomms7925

Kashtan N, Roggensack SE, Rodrigue S, Thompson JW, Biller SJ, Coe A, Ding H, Marttinen P, Malmstrom RR, Stocker R, Follows MJ, Stepanauskas R, Chisholm SW. 2014. Single-cell genomics reveals hundreds of coexisting subpopulations in wild Prochlorococcus. Science 344:416–20. doi:10.1126/science.1248575

Katoh K, Misawa K, Kuma KI, Miyata T. 2002. MAFFT: A novel method for rapid multiple sequence alignment based on fast Fourier transform. Nucleic Acids Res 30:3059–3066. doi:10.1093/nar/gkf436

Kawachi M, Inouye I, Honda D, O’Kelly CJ, Bailey JC, Bidigare RR, Andersen RA. 2002. The Pinguiophyceae classis nova, a new class of photosynthetic stramenopiles whose members produce large amounts of omega-3 fatty acids. Phycol Res 50:31–47. doi: 10.1046/j.1440-1835.2002.00260.x

Kojima K, Suzuki-Maenaka T, Kikuchi T, Nakamoto H. 2006. Roles of the cyanobacterial isiABC operon in protection from oxidative and heat stresses. Physiol Plant 128:507–519. doi:10.1111/j.1399-3054.2006.00781.x

Lambert BS, Groussman RD, Schatz MJ, Coesel SN, Durham BP, Alverson AJ, White AE, Armbrust EV. 2022. The dynamic trophic architecture of open-ocean protist communities revealed through machine-guided metatranscriptomics. Proc Natl Acad Sci 119. doi:10.1073/pnas.2100916119

Lasek-Nesselquist E, Johnson MD. 2019. A Phylogenomic Approach to Clarifying the Relationship of Mesodinium within the Ciliophora: A Case Study in the Complexity of Mixed-Species Transcriptome Analyses. Genome Biol Evol 11:3218–3232. doi: 10.1093/gbe/evz233

Letunic I, Bork P. 2021. Interactive tree of life (iTOL) v5: An online tool for phylogenetic tree display and annotation. Nucleic Acids Res 49:W293–W296. doi: 10.1093/nar/gkab301

Lommer M, Roy A-S, Schilhabel M, Schreiber S, Rosenstiel P, LaRoche J. 2010. Recent transfer of an iron-regulated gene from the plastid to the nuclear genome in an oceanic diatom adapted to chronic iron limitation. BMC Genomics 11:718. doi:10.1186/1471-2164-11-718

Lommer M, Specht M, Roy AS, Kraemer L, Andreson R, Gutowska MA, Wolf J, Bergner S V, Schilhabel MB, Klostermeier UC, Beiko RG, Rosenstiel P, Hippler M, Laroche J. 2012. Genome and low-iron response of an oceanic diatom adapted to chronic iron limitation. Genome Biol 13:R66. doi:10.1186/gb-2012-13-7-r66

Mackey KRM, Post AF, McIlvin MR, Cutter G a., John SG, Saito M a. 2015. Divergent responses of Atlantic coastal and oceanic Synechococcus to iron limitation. Proc Natl Acad Sci 201509448. doi:10.1073/pnas.1509448112

Marchetti A, Schruth DM, Durkin CA, Parker MS, Kodner RB, Berthiaume CT, Morales R, Allen AE, Armbrust EV. 2012. Comparative metatranscriptomics identifies molecular bases for the physiological responses of phytoplankton to varying iron availability. Proc Natl Acad Sci 109:E317–25. doi:10.1073/pnas.1118408109

Mayta ML, Arce RC, Zurbriggen MD, Valle EM, Hajirezaei MR, Zanor MI, Carrillo N. 2019. Expression of a Chloroplast-Targeted Cyanobacterial Flavodoxin in Tomato Plants Increases Harvest Index by Altering Plant Size and Productivity. Front Plant Sci 10:1–13. doi:10.3389/fpls.2019.01432

Mock T, Otillar RP, Strauss J, McMullan M, Paajanen P, Schmutz J, Salamov A, Sanges R, Toseland A, Ward BJ, Allen AE, Dupont CL, Frickenhaus S, Maumus F, Veluchamy A, Wu T, Barry KW, Falciatore A, Ferrante MI, Fortunato AE, Glöckner G, Gruber A, Hipkin R, Janech MG, Kroth PG, Leese F, Lindquist EA, Lyon BR, Martin J, Mayer C, Parker M, Quesneville H, Raymond JA, Uhlig C, Valas RE, Valentin KU, Worden AZ, Armbrust EV, Clark MD, Bowler C, Green BR, Moulton V, van Oosterhout C, Grigoriev I V. 2017. Evolutionary genomics of the cold-adapted diatom Fragilariopsis cylindrus. Nature. doi:10.1038/nature20803

Mondal J, Bruce BD. 2018. Ferredoxin: the central hub connecting photosystem I to cellular metabolism. Photosynthetica 56:279–293. doi:10.1007/s11099-018-0793-9

Moore CM, Mills MM, Arrigo KR, Berman-Frank I, Bopp L, Boyd PW, Galbraith ED, Geider RJ, Guieu C, Jaccard SL, Jickells TD, La Roche J, Lenton TM, Mahowald NM, Marañón E, Marinov I, Moore JK, Nakatsuka T, Oschlies A, Saito M a., Thingstad TF, Tsuda A, Ulloa O. 2013. Processes and patterns of oceanic nutrient limitation. Nat Geosci 6:701–710. doi:10.1038/ngeo1765

Nam O, Grouneva I, Mackinder LCM. 2022. Endogenous GFP tagging in the diatom Thalassiosira pseudonana. bioRxiv 1–23. doi:10.1101/2022.09.30.510313

Nekrasov V, Staskawicz B, Weigel D, Jones JDG, Kamoun S. 2013. Targeted mutagenesis in the model plant Nicotiana benthamiana using Cas9 RNA-guided endonuclease. Nat Biotechnol 31:688–691. doi:10.1038/nbt.2654

Oudot-Le Secq MP, Grimwood J, Shapiro H, Armbrust EV, Bowler C, Green BR. 2007. Chloroplast genomes of the diatoms Phaeodactylum tricornutum and Thalassiosira pseudonana: Comparison with other plastid genomes of the red lineage. Mol Genet Genomics 277:427–439. doi:10.1007/s00438-006-0199-4

Park J, Durham BP, Key RS, Groussman RD, Pinedo-Gonzalez P, Hawco NJ, John SG, Carlson MCG, Lindell D, Juranek L, Ferrón S, Ribalet F, Armbrust EV, Ingalls AE, Bundy RM. 2022. Siderophore production and utilization by microbes in the North Pacific Ocean. bioRxiv 2022.02.26.482025. doi:/10.1101/2022.02.26.482025

Pierella Karlusich JJ, Carrillo N. 2017. Evolution of the acceptor side of photosystem I: ferredoxin, flavodoxin, and ferredoxin-NADP + oxidoreductase. Photosynth Res 134:235–250. doi:10.1007/s11120-017-0338-2

Pierella Karlusich JJ, Ceccoli RD, Graña M, Romero H, Carrillo N, Grana M, Romero H, Carrillo N. 2015. Environmental selection pressures related to iron utilization are involved in the loss of the flavodoxin gene from the plant genome. Genome Biol Evol 7:750–767. doi: 10.1093/gbe/evv031

Pinedo-González P, Hawco NJ, Bundy RM, Armbrust EV, Follows MJ, Cael BB, White AE, Ferrón S, Karl DM, John SG. 2020. Anthropogenic Asian aerosols provide Fe to the North Pacific Ocean. Proc Natl Acad Sci U S A 117:27862–27868. doi:10.1073/pnas.2010315117

Price NM, Harrison GI, Hering JG, Hudson RJ, Nirel PM, Palenik B, Morel FM. 1989. Preparation and chemistry of the artificial algal culture medium Aquil. Biol Oceanogr 6:443–461.

Robinson MD, McCarthy DJ, Smyth GK. 2009. edgeR: A Bioconductor package for differential expression analysis of digital gene expression data. Bioinformatics 26:139–140. doi:10.1093/bioinformatics/btp616

Roy A-S, Woehle C, LaRoche J. 2020. The Transfer of the Ferredoxin Gene From the Chloroplast to the Nuclear Genome Is Ancient Within the Paraphyletic Genus Thalassiosira. Front Microbiol 11:1–15. doi:10.3389/fmicb.2020.523689

Singh AK, Elvitigala T, Cameron JC, Ghosh BK, Bhattacharyya-Pakrasi M, Pakrasi HB. 2010. Integrative analysis of large scale expression profiles reveals core transcriptional response and coordination between multiple cellular processes in a cyanobacterium. BMC Syst Biol 4. doi:10.1186/1752-0509-4-105

Singh AK, Li H, Sherman LA. 2004. Microarray analysis and redox control of gene expression in the cyanobacterium Synechocystis sp. PCC 6803. Physiol Plant 120:27–35. doi:10.1111/j.0031-9317.2004.0232.x

Smillie RM. 1965. Isolation of two proteins with chloroplast ferredoxin activity from a blue-green alga. Biochem Biophys Res Commun 20:621–629. doi: 10.1016/0006-291X(65)90445-6

Smith SR, Gillard JTF, Kustka AB, McCrow JP, Badger JH, Zheng H, New AM, Dupont CL, Obata T, Fernie AR, Allen AE. 2016. Transcriptional Orchestration of the Global Cellular Response of a Model Pennate Diatom to Diel Light Cycling under Iron Limitation. PLoS Genet 12:e1006490. doi:10.1371/journal.pgen.1006490

Stamatakis A. 2014. RAxML version 8: A tool for phylogenetic analysis and post-analysis of large phylogenies. Bioinformatics 30:1312–1313. doi:10.1093/bioinformatics/btu033

Strand TA, Lale R, Degnes KF, Lando M, Valla S. 2014. A new and improved host-independent plasmid system for RK2-based conjugal transfer. PLoS One 9:1–6. doi: 10.1371/journal.pone.0090372

Thaben PF, Westermark PO. 2014. Detecting rhythms in time series with RAIN. J Biol Rhythms 29:391–400. doi: 10.1177/0748730414553029

Thompson AW, Huang K, Saito MA, Chisholm SW. 2011. Transcriptome response of high- and low-light-adapted Prochlorococcus strains to changing iron availability. ISME J 5:1580–1594. doi:10.1038/ismej.2011.49

Tognetti VB, Palatnik JF, Fillat MF, Melzer M, Hajirezaei M, Valle EM, Carrillo N. 2006. Functional replacement of ferredoxin by a cyanobacterial flavodoxin in Tobacco confers broad-range stress tolerance. Plant Cell 18:2035–2050. doi:10.1105/tpc.106.042424.1

Villareal TA, Brown CG, Brzezinski MA, Krause JW, Wilson C. 2012. Summer diatom blooms in the north Pacific subtropical gyre: 2008-2009. PLoS One 7:2008–2009. doi:10.1371/journal.pone.0033109

Wagih O. 2017. Ggseqlogo: A versatile R package for drawing sequence logos. Bioinformatics 33:3645–3647. doi: 10.1093/bioinformatics/btx469

Weber E, Engler C, Gruetzner R, Werner S, Marillonnet S. 2011. A modular cloning system for standardized assembly of multigene constructs. PLoS One 6:e16765. doi:10.1371/journal.pone.0016765

WG Zumft, Spiller H. 1971. Characterization of a flavodoxin from the green alga Chlorella. Biochem Biophys Res Commun 44:995–1000. doi:10.1016/0006-291X(71)90057-X

Whitney LAP, Lins JJ, Hughes MP, Wells ML, Dreux Chappell P, Jenkins BD. 2011. Characterization of putative iron responsive genes as species-specific indicators of iron stress in Thalassiosiroid diatoms. Front Microbiol 2:1–14. doi:10.3389/fmicb.2011.00234

Wilson ST, Aylward FO, Ribalet F, Barone B, Casey JR, Connell PE, Eppley JM, Ferron S, Fitzsimmons JN, Hayes CT, Romano AE, Turk-Kubo KA, Vislova A, Virginia Armbrust E, Caron DA, Church MJ, Zehr JP, Karl DM, De Long EF. 2017. Coordinated regulation of growth, activity and transcription in natural populations of the unicellular nitrogen-fixing cyanobacterium Crocosphaera. Nat Microbiol 2:1–9. doi:10.1038/nmicrobiol.2017.118

Wu M, McCain JSP, Rowland E, Middag R, Sandgren M, Allen AE, Bertrand EM. 2019. Manganese and iron deficiency in Southern Ocean Phaeocystis antarctica populations revealed through taxon-specific protein indicators. Nat Commun 10:3582. doi:10.1038/s41467-019-11426-z

Yamada N, Bolton JJ, Trobajo R, Mann DG, Dąbek P, Witkowski A, Onuma R, Horiguchi T, Kroth PG. 2019. Discovery of a kleptoplastic ‘dinotom’ dinoflagellate and the unique nuclear dynamics of converting kleptoplastids to permanent plastids. Sci Rep 9:1–13. doi: 10.1038/s41598-019-46852-y

Yoch DC, Valentine RC. 1972. Ferredoxins and flavodoxins of bacteria. Annu Rev Microbiol 26:139–162. doi:10.1146/annurev.mi.26.100172.001035

Yousef N, Pistorius EK, Michel KP. 2003. Comparative analysis of idiA and isiA transcription under iron starvation and oxidative stress in Synechococcus elongatus PCC 7942 wild-type and selected mutants. Arch Microbiol 180:471–483. doi:10.1007/s00203-003-0618-4

Zheng M, Doan B, Schneider TD, Storz G. 1999. OxyR and SoxRS regulation of fur. J Bacteriol 181:4639–4643. doi:10.1128/jb.181.15.4639-4643.1999

Zurbriggen MD, Tognetti VB, Fillat MF, Hajirezaei MR, Valle EM, Carrillo N. 2008. Combating stress with flavodoxin: a promising route for crop improvement. Trends Biotechnol 26:531–537. doi:10.1016/j.tibtech.2008.07.001

