## Supplemental Figures for "Divergent functions of two clades of flavodoxin in diatoms mitigate oxidative stress and iron limitation"

**Figure S1.** Flavodoxins in publicly available databases

**A.**

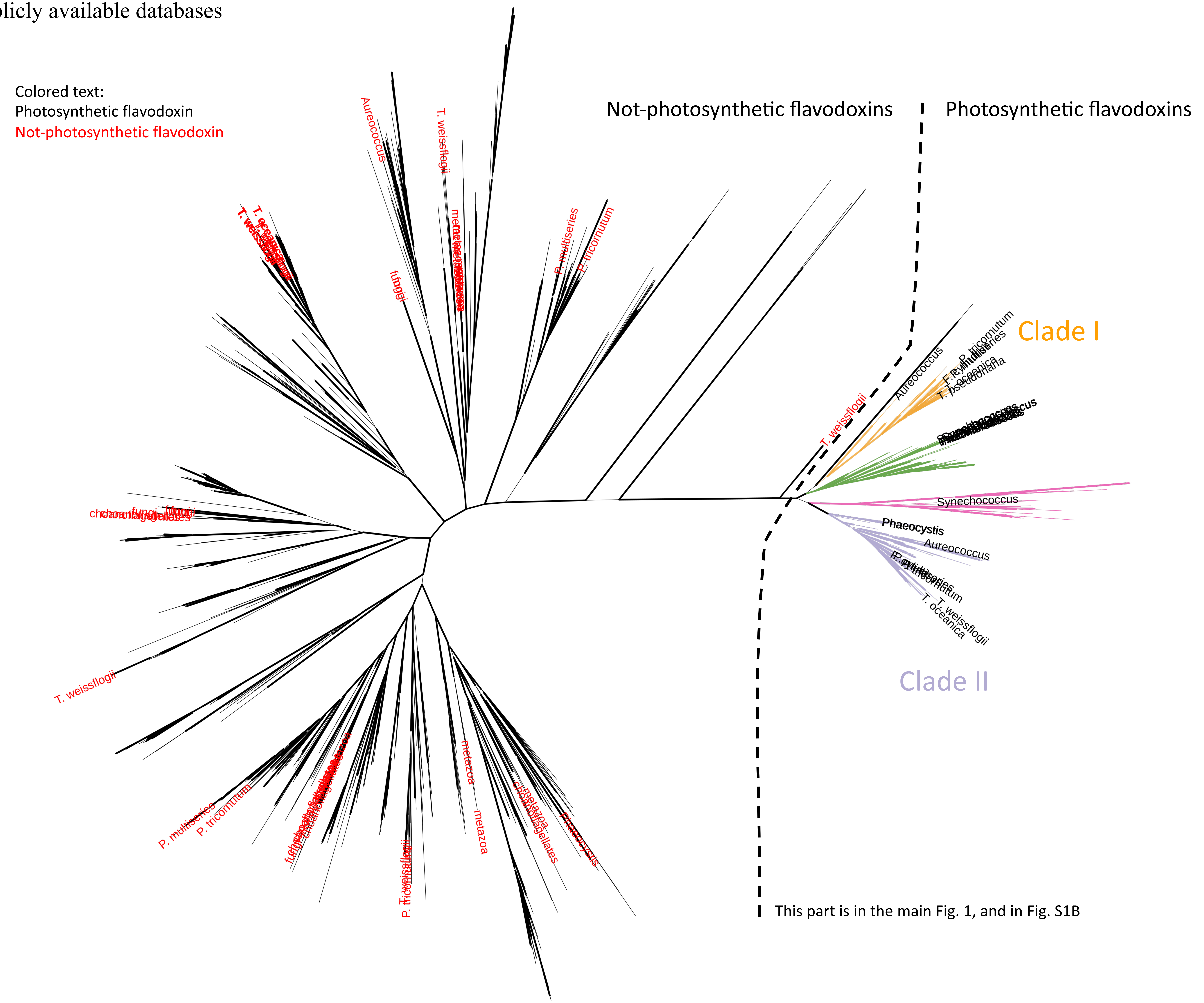

**Figure S1.** Flavodoxins in publicly available databases

**B.**

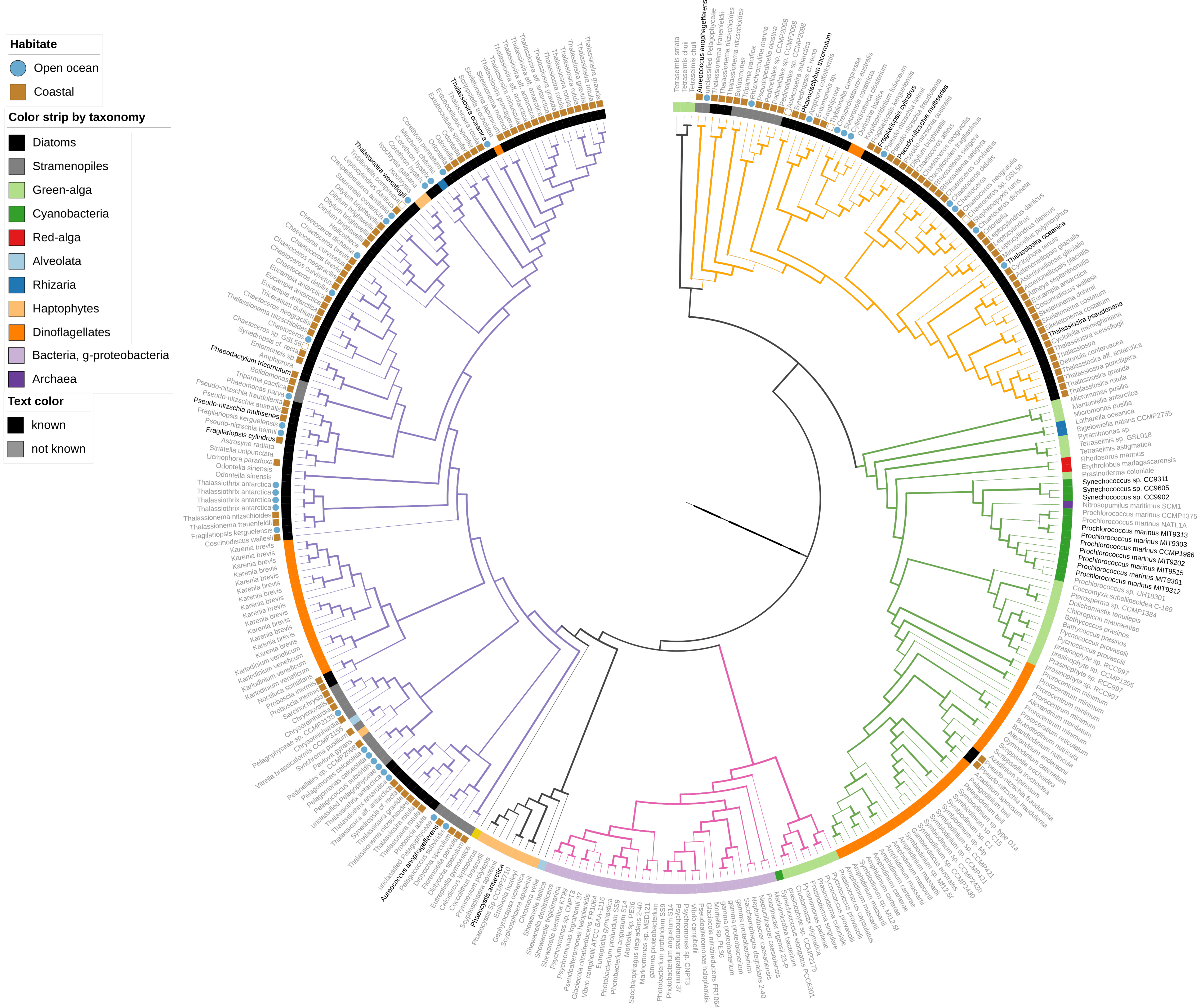

**Figure S1.** Flavodoxins in publicly available databases

**C.**

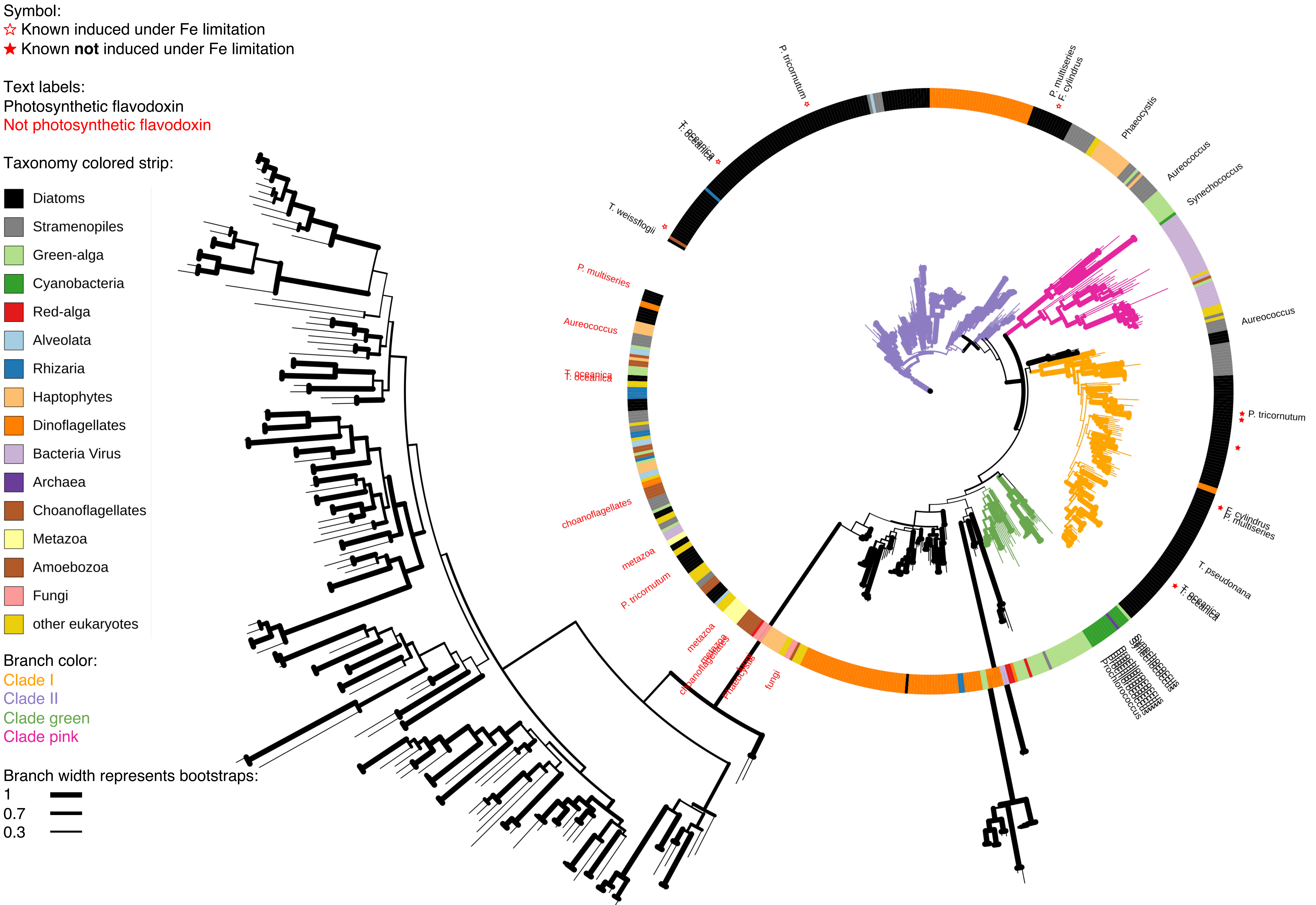

**Figure S1.** Flavodoxins in publicly available databases

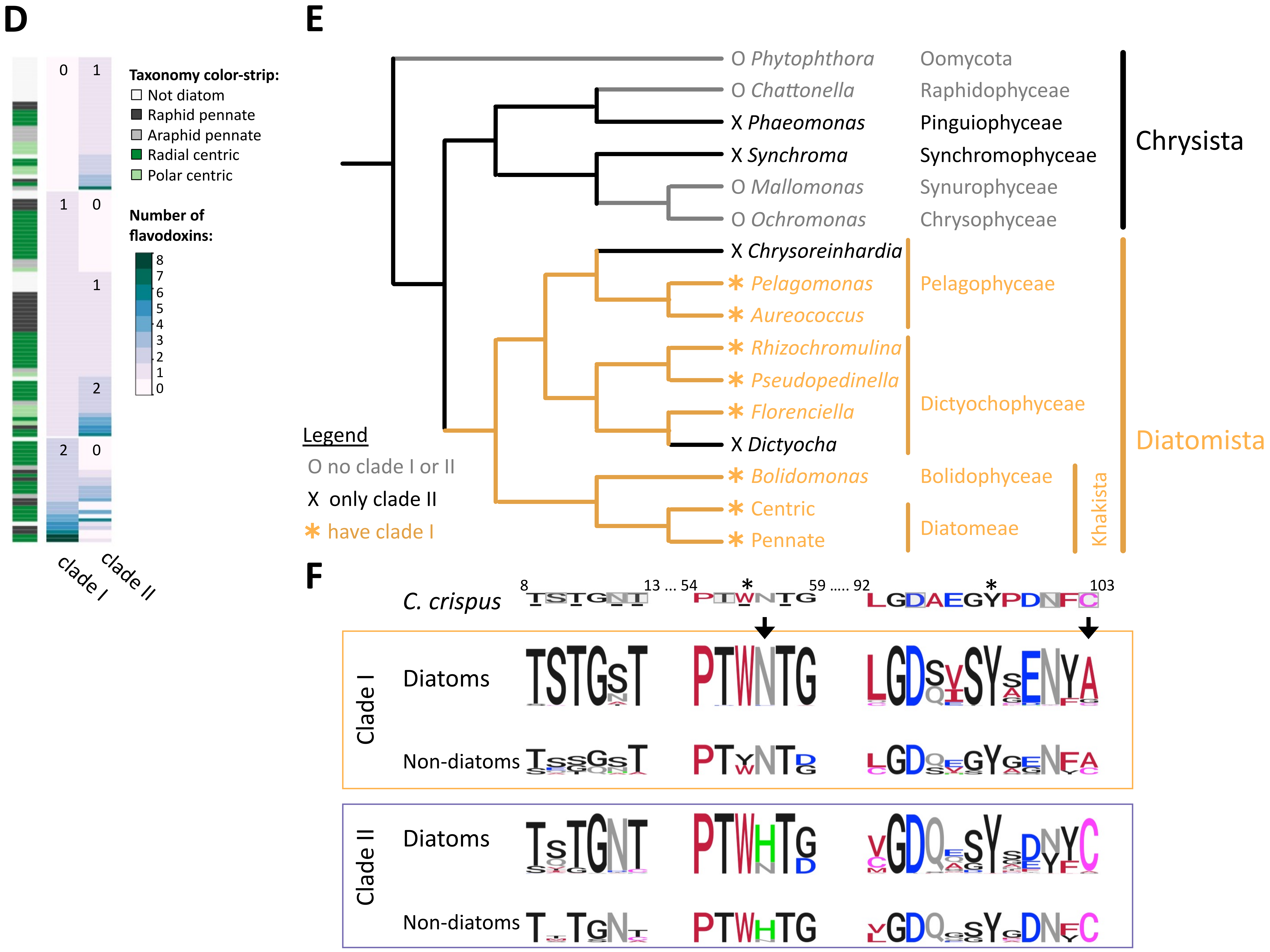

**Figure S2.** Diatoms cultures transcriptomes

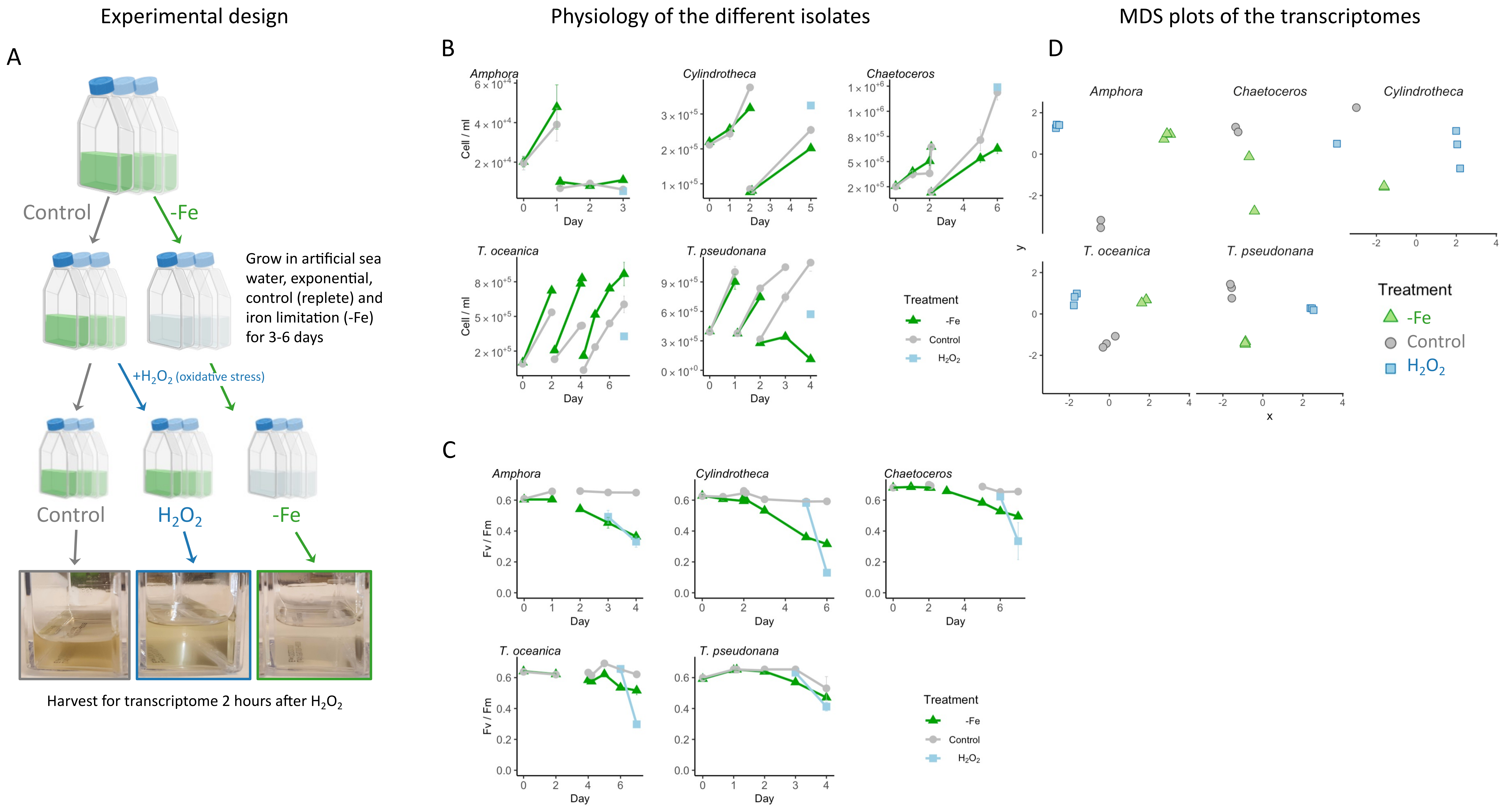

**Figure S3.** Flavodoxin knock-out in *T. pseudonana*

**A.**

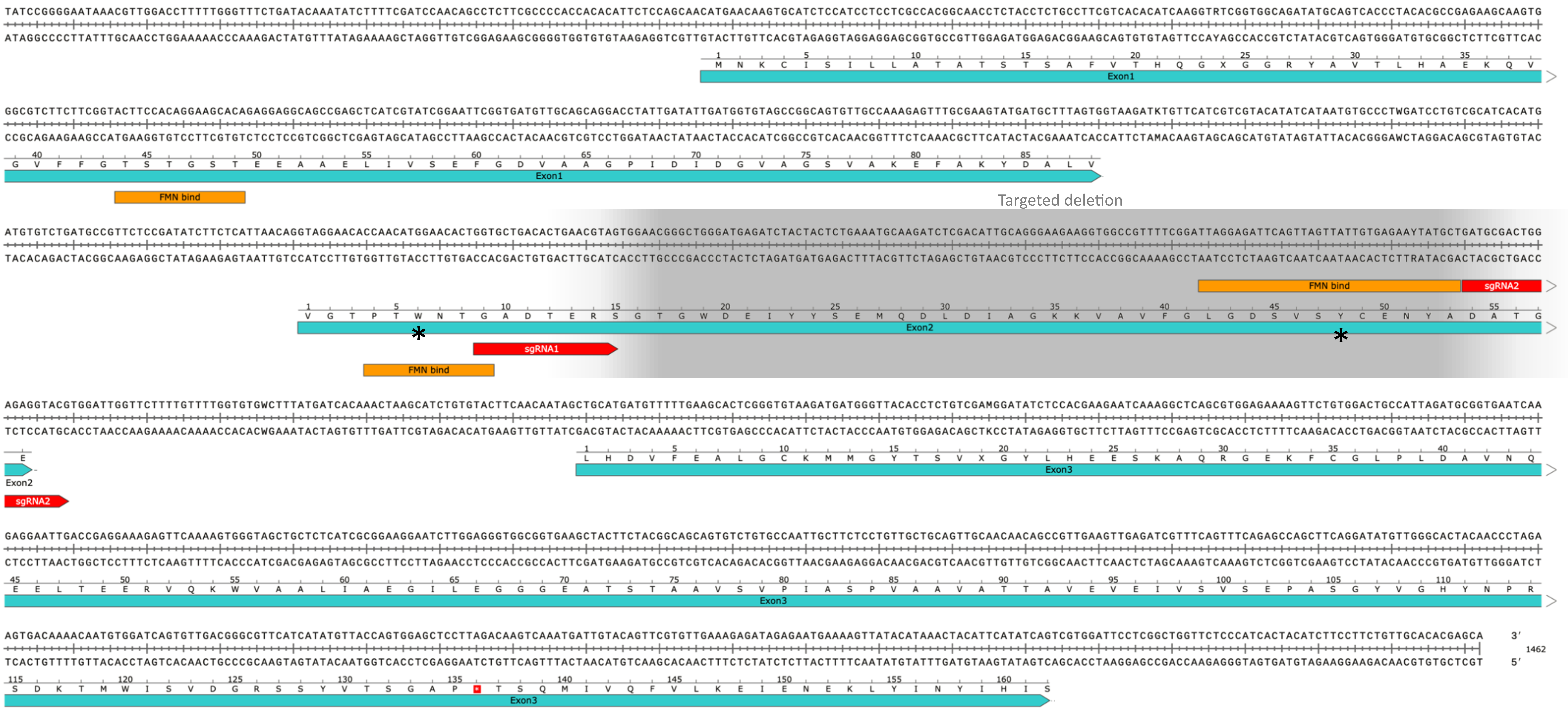

**Figure S3.** Flavodoxin knock-out in *T. pseudonana*

**B**

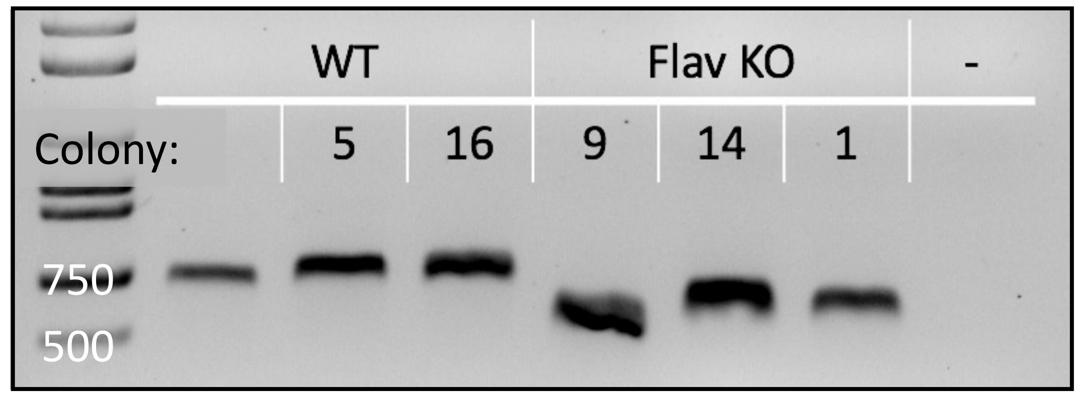


**C**

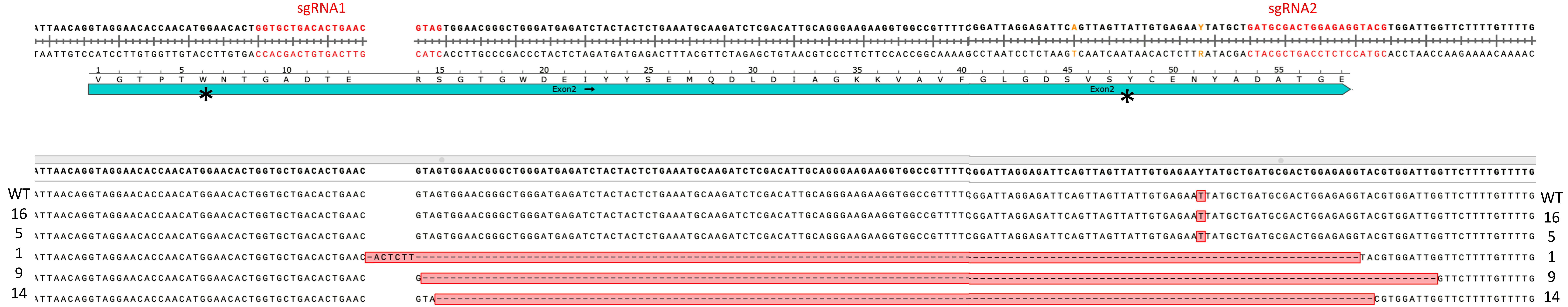

**Figure S3.** Flavodoxin knock-out in *T. pseudonana*

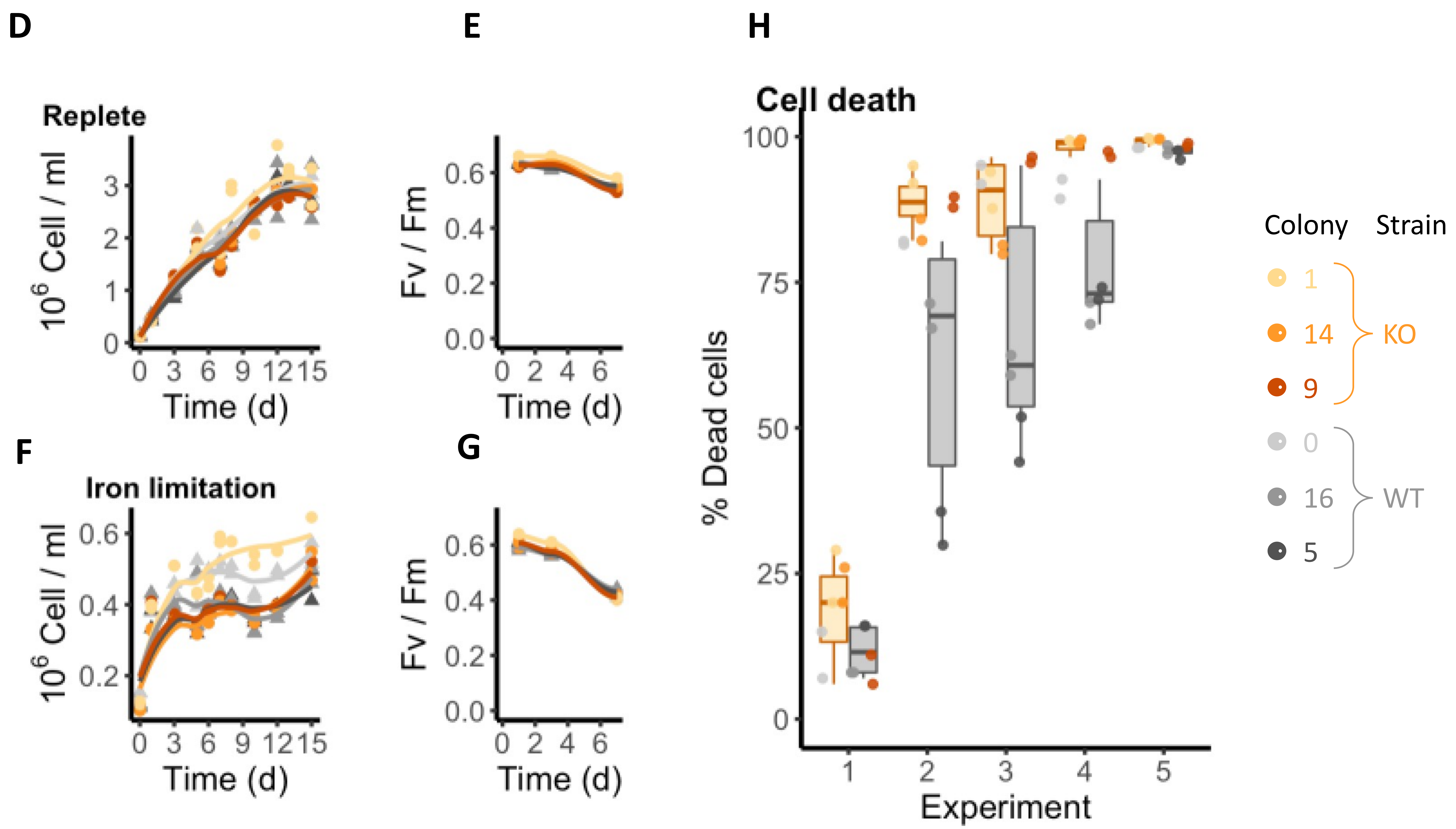

**Figure S4.** Flavodoxins in the North Pacific Subtropical Gyre

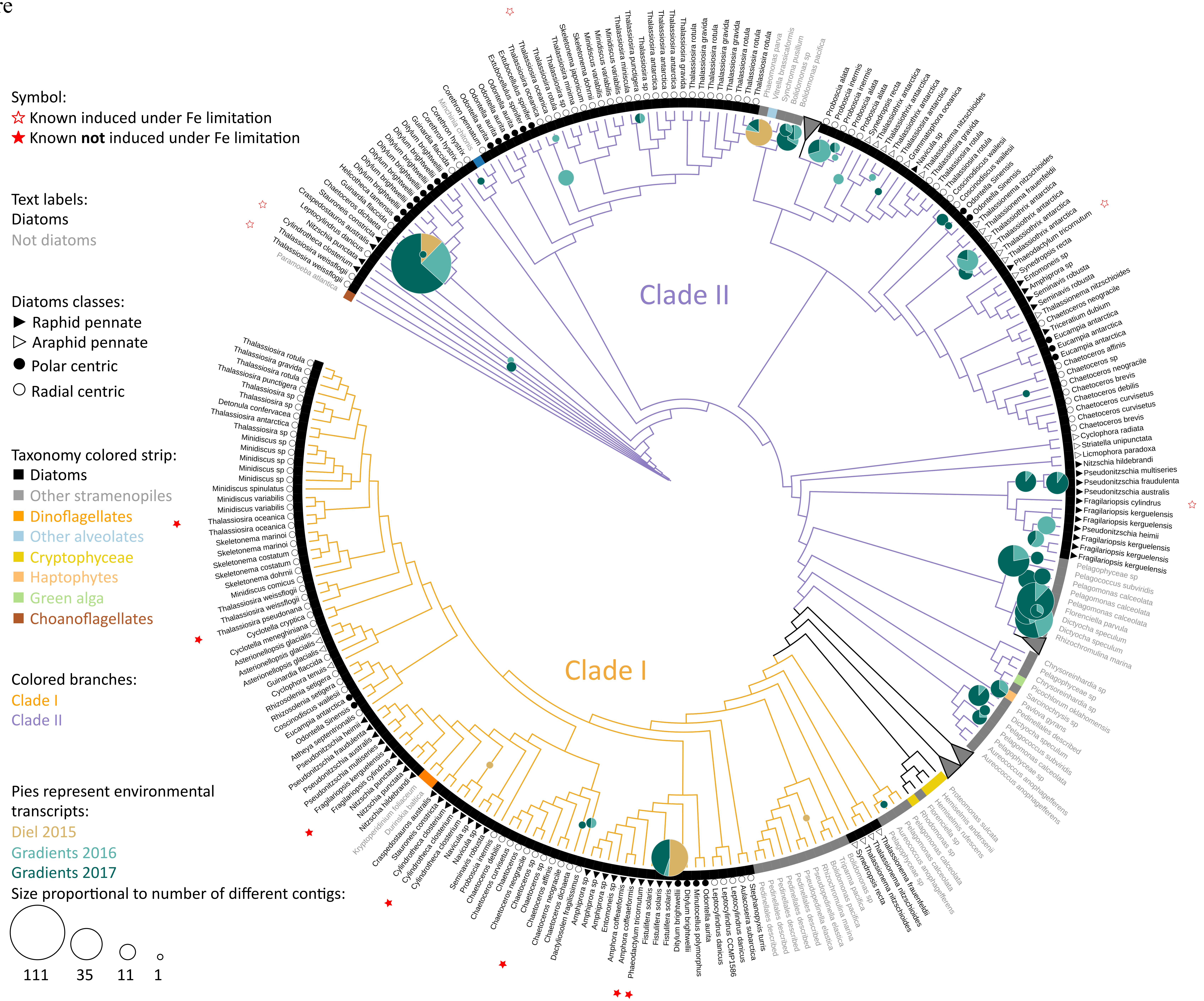

**Figure S5.** Transcriptional patterns of flavodoxin genes in the North Pacific

**A.**

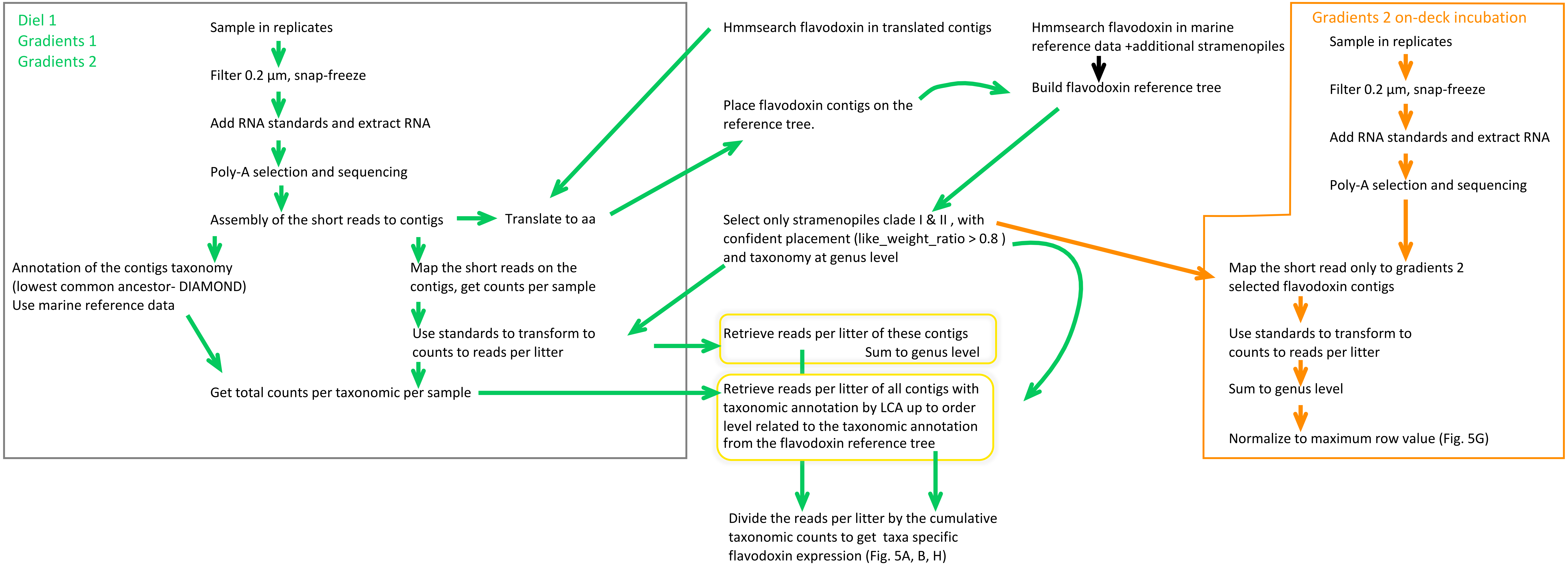

**Figure S5.** Transcriptional patterns of flavodoxin genes in the North Pacific

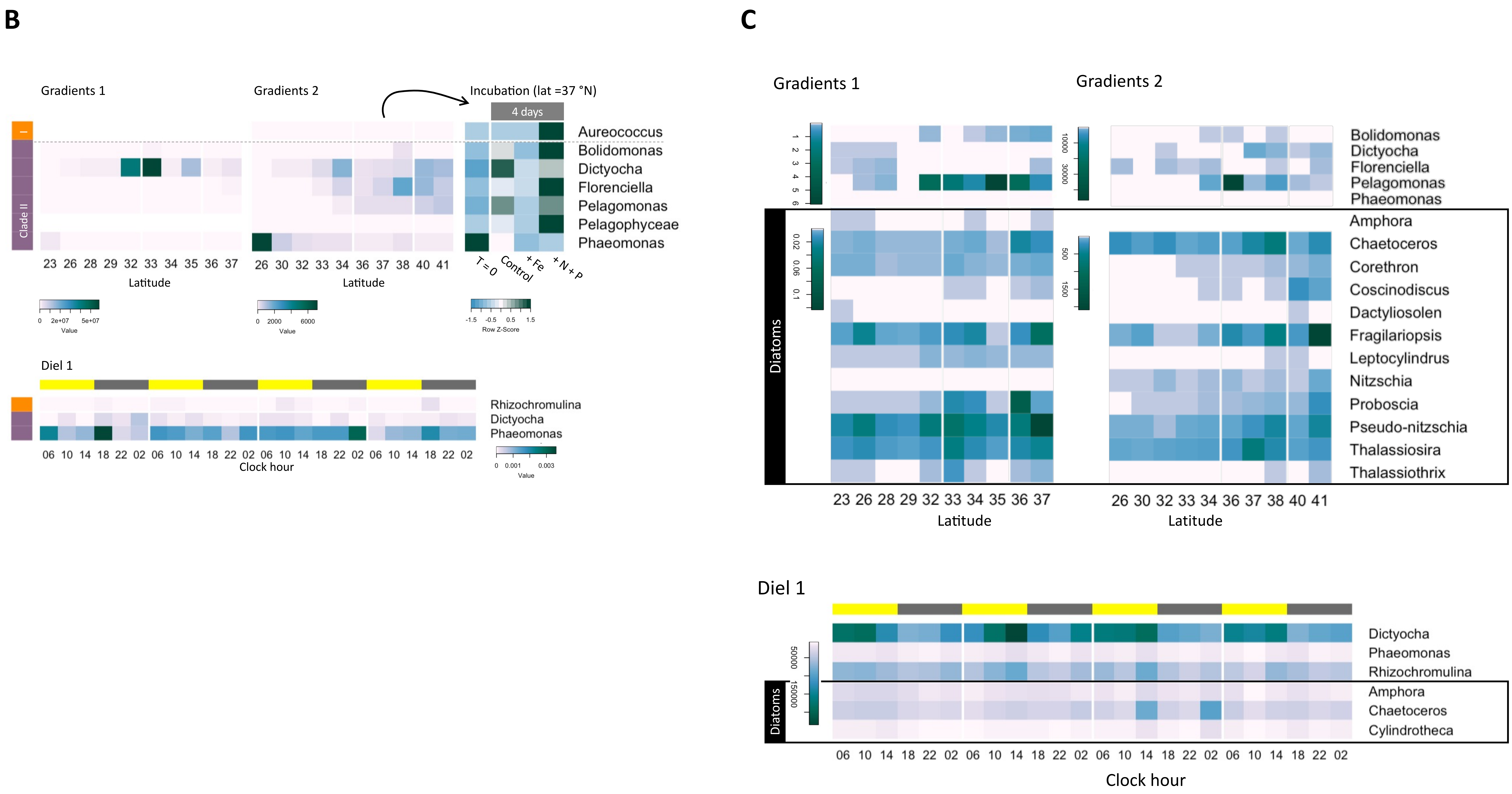
